# Maintaining hypoxia environment of subchondral bone alleviates osteoarthritis progression

**DOI:** 10.1101/2022.03.17.484053

**Authors:** Hao Zhang, Lipeng Wang, Jin Cui, Sicheng Wang, Yafei Han, Hongda Shao, Cheng Wang, Yan Hu, Xiaoqun Li, Qirong Zhou, Jiawei Guo, Xinchen Zhuang, Shihao Sheng, Tao Zhang, Dongyang Zhou, Jiao Chen, Fuxiao Wang, Qianmin Gao, Yingying Jing, Xiao Chen, Jiacan Su

## Abstract

Abnormal subchondral bone remodeling featured by over-activated osteoclastogenesis leads to articular cartilage degeneration and osteoarthritis (OA) progression, but the mechanism is still unclear. In this study, we used lymphocyte cytosolic protein 1 (*Lcp1*) knock-out mice to suppress subchondral osteoclast formation in mice OA model with anterior cruciate ligament transection (ACLT) and *Lcp1*^-/-^ mice showed decreased bone remodeling and sensory innervation in subchondral bone accompanied by retarded cartilage degeneration. For mechanisms, in wildtype mice with ACLT the activated osteoclasts in subchondral bone induced type-H vessels and elevated oxygen concentration which ubiquitylated hypoxia-inducible factor 1α (HIF-1α), vital for maintaining chondrocyte homeostasis in articular chondrocytes and led to cartilage degeneration. Deletion of *Lcp1* impeded osteoclast-mediated angiogenesis, which maintained the low levels of oxygen partial pressure (pO_2_) in subchondral bone as well as the whole joint and delayed the OA progression. Stabilization of HIF-1α delayed cartilage degeneration and knockdown of *Hif1a* abolished the protective effects of *Lcp1* knockout. Notably, we identified a novel subgroup of hypertrophic chondrocytes highly associated with OA by single cell sequencing analysis of human articular chondrocytes. Lastly, we showed that Oroxylin A, an *Lcp1-*encoded protein L-plastin (LPL) inhibitor, could alleviate OA progression. In conclusion, maintaining hypoxic environment in subchondral bone is an attractive strategy for OA treatment.

**Teaser:** Inhibiting subchondral osteoclastogenesis alleviates OA progression via maintaining joint hypoxia environment.

## INTRODUCTION

OA is a complex disease affecting the whole joint, characterized by cartilage degeneration, aberrant bone remodeling, osteophyte formation and joint inflammation (*1*). As the leading cause of disability and pain, OA affects over 300 million people worldwide (*2, 3*). The current treatment algorithm including self-education and cycloxygenase-2 (COX-2) inhibitors mainly helps symptom alleviation and no disease-modifying osteoarthritis drug (DMOAD) is available due to the limited understanding of OA pathogenesis (*4*).

OA is featured by subchondral bone changes in clinical findings (*5*). Subchondral bone subjacent to cartilage provides nutritional and mechanical support for cartilage (*6*). Subchondral bone marrow edema, formation of osteocysts and sclerosis could be found in most OA patients. Subchondral bone marrow edema first appears in MRI images followed by osteocyst formation in early OA patients and the subchondral bone marrow edema area in MRI corresponds to the degenerated cartilage above (*7, 8*). The roles of subchondral bone in OA progression remain unclear.

Over-activated osteoclasts in subchondral bone are closely associated with OA progression (*9*). Physiologically the bone remodeling activity and number of osteoclasts are strictly controlled in the subchondral bone and the number of osteoclasts dramatically increases at the early stage of OA mice model (*10, 11*). Several hypotheses regarding osteoclast roles in OA have been proposed. Osteoclast precursors migrate into the cartilage layer and directly contact with hypertrophic chondrocytes to degrade the osteochondral junction and articular cartilage (*12*). Growth factors released from the bone matrix through osteoclastic bone resorption including transforming growth factor beta 1 (TGF-β1), insulin-like growth factor 1 (IGF-1), platelet-derived growth factor BB (PDGF-BB) regulate chondrocyte metabolism (*13, 14*). Nevertheless, the number of osteoclasts significantly drops after reaching the peak in OA model, the cartilage degeneration continuously deteriorates. Thus, the roles of osteoclasts in crosstalk between subchondral bone and chondrocytes remain mysterious.

Due to lack of blood vessels, subchondral bone and cartilage remain hypoxic, which is vital for chondrocyte homeostasis (*15*). Hypervascularization in subchondral bone is the hallmark and drug target of OA progression. Angiogenesis stimulated by elevated PDGF-BB in subchondral bone contributes to osteoarthritis development (*13*). Administration of bevacizumab, a vascular endothelial growth factor (VEGF) blocker, inhibits angiogenesis and mitigates OA (*13*). Thus, we hypothesize that blood vessels formation induced by osteoclasts in subchondral bone in early stage of OA alters the joint hypoxia environment and contributes to sustained cartilage degeneration.

In this study, we used *Lcp1* knockout mice with impaired osteoclast formation as we previously reported and established OA model with ACLT (*18*). *Lcp1*^-/-^ mice after ACLT showed preserved articular cartilage and delayed OA progression. Mechanistically, angiogenesis by osteoclast activation elevated the concentration of O_2_ in subchondral bone and cartilage. The disrupted joint hypoxia environment with elevated oxygen partial pressure promoted chondrocytes degeneration by abolishing HIF-1α functions and stabilizing Hif-1α functions prevented cartilage destruction.

## RESULTS

### L-plastin upregulated in subchondral bone correlates with increased osteoclast activity in early OA

To investigate the involvement of osteoclasts and LPL in OA, we analyzed the Micro-CT and knee section of wildtype (WT) mice after ACLT at different time points. During OA progression, the thickness of hyaline cartilage (HC) decreased while the thickness of calcified cartilage (CC) gradually duplicated and ratio of HC/CC decreased to average 0.81 at 8 weeks (Fig.1, A and B). The cartilage degeneration kept progressing during 8 weeks with the arising Osteoarthritis Research Society International (OARSI) grade (Fig.1, C and D). The bone mass of subchondral bone decreased at 2 weeks after ACLT with bone volume/total volume (BV/TV) ranging from 34.3%-55.8% but BV/TV increased to 56.6%-76.2% at 4 and 8 weeks after ACLT (Fig.1, E to G). Correspondingly, the number of tartrate resistant acid phosphatase (TRAP) positive cells increased rapidly during the first two weeks after ACLT to the peak of average 13.4 cells per mm^2^ and decreased after 2 weeks (Fig.1, H and I). LPL is exclusively expressed in myeloid lineage cells, and vital for osteoclast fusion and mature osteoclast formation (*18*). Herein, the immunohistochemistry (IHC) results showed that the number of LPL^+^ cells in subchondral bone was consistent with that of TRAP^+^ cells, reaching the peak of average 10.8 cells/mm^2^ (Fig.1, J and K). No LPL^+^ cell was observed in cartilage, few were found surrounding subchondral trabecular bone and abundant were detected around the primary spongiosa near epiphysis (fig.S1A). Two weeks after ACLT, a significantly increased number of LPL^+^ cells were observed around the subchondral trabecular bone, but not in cartilage (fig.S1B). Taken together, under physiological condition, nearly no LPL^+^ or TRAP^+^ cells could be detected in subchondral bone, while 2 weeks after ACLT the number of LPL^+^ and TRAP^+^ cells is significantly increased, indicating an accelerated bone remodeling in subchondral bone in early stage of OA.

**Fig.1.**
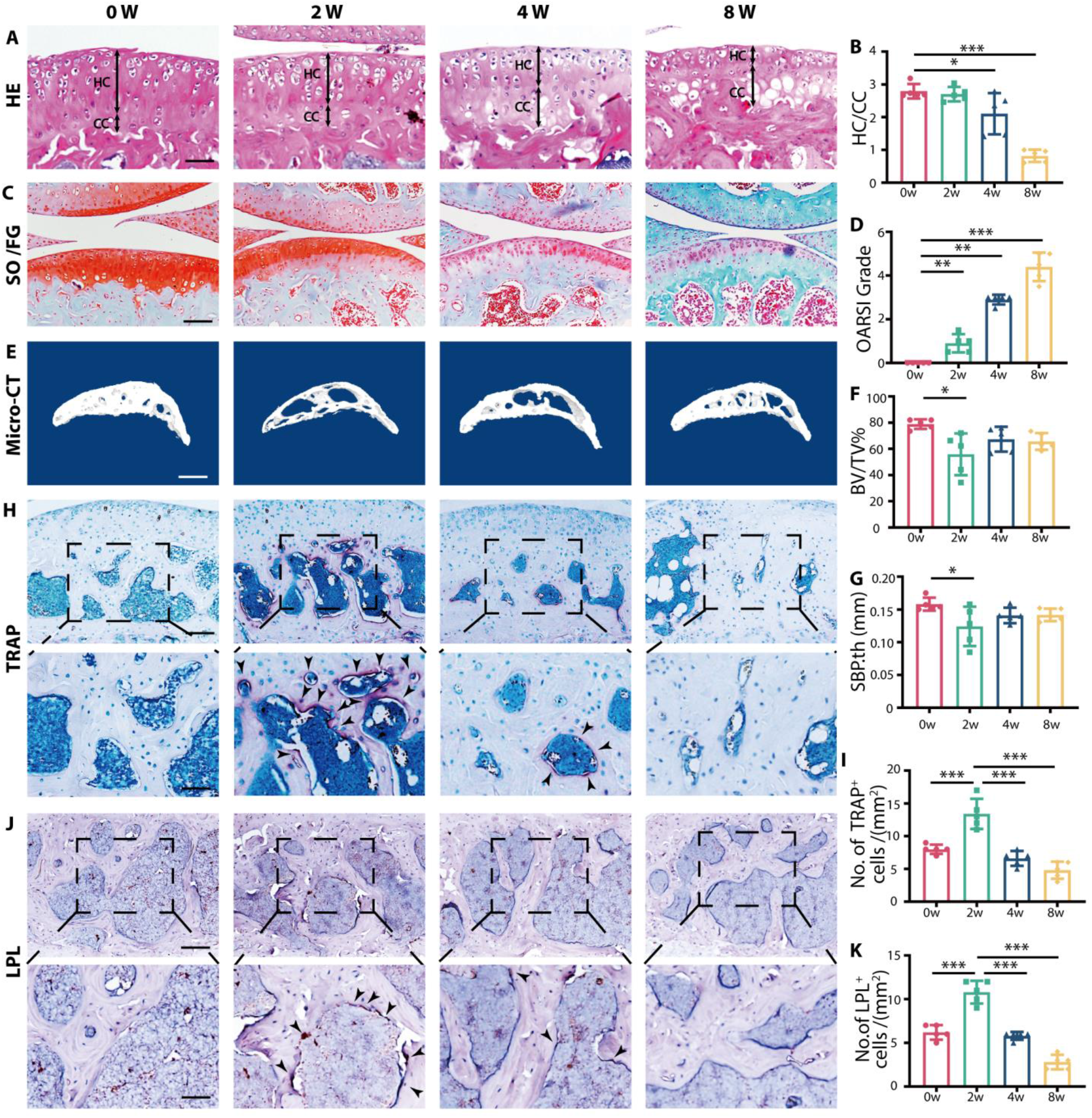
L-plastin upregulated in subchondral bone correlates with increased osteoclast activity in early stage of OA. **(A)** Representative images of Hematoxylin and eosin staining of proximal tibia articular cartilage at 0,2,4 and 8 weeks after ACLT. Double-headed arrows label range of HC and CC. Scale bar, 50μm. **(B)** Ratio of HC and CC thickness quantitative analysis. **(C)** Safranin O/Fast Green staining of knee articular cartilage at 0,2,4 and 8 weeks after operation. Scale bar 100μm. **(D)** OARSI grade of knee articular cartilage. **(E)** Three-dimensional images of the sagittal plane of medial tibial subchondral bone at 0,2,4 and 8 weeks after ACLT. Scale bar, 500μm. **(F-G)** Micro-CT quantitative analysis of tibial subchondral bone, bone volume/tissue volume (BV/TV, %) (F), and subchondral bone plate thickness (SBP. th, mm) (G). **(H)** TRAP staining image of tibial subchondral bone at 0,2,4 and 8 weeks after ACLT. Scale bar, 100 μm (top). Scale bar, 50 μm (bottom) **(I)** Quantitative analysis of TRAP-positive cells in subchondral bone marrow. **(J)** Representative images of LPL protein immunohistochemistry in tibial subchondral bone at 0,2,4 and 8 weeks after ACLT. Scale bar =100 μm (top). Scale bar, 50 μm (bottom) **(K)** Quantitative analysis of LPL positive osteoclast in subchondral bone marrow. Statistical analysis was performed using one-way ANOVA analysis; n=5 per group; Data are presented as means ± SEM. *P < 0.05, **P < 0.01, and ***P < 0.001.

### *Lcp1* knockout reduces subchondral bone resorption and ameliorates articular cartilage degeneration

To explore the role of LPL in OA progression, we generated *Lcp1*^-/-^ mice. Micro-CT results showed that the BV/TV in subchondral bone marrow of *Lcp1* deletion mice significantly increased to average 75.1% and 81.9% compared to their littermate average 55.8% and 67.4% at 2 and 4 weeks (Fig.2, A and B). There was no statistical difference of subchondral bone plate thickness between the two groups (Fig.2C). As our previous study showed, the number of TRAP^+^ cells in subchondral bone marrow in *Lcp1* deletion mice was almost half of numbers (53.7% and 54.2%) in WT group at 2 and 4 weeks after ACLT (Fig.2, D and E). The results of Safranin O/Fast green staining showed that the OARSI grade increased at 4 and 8 weeks after ACLT in WT mice. *Lcp1* knockout significantly decreased 1.25 and 2.25 grade compared with WT mice (Fig.2, F and G). The ratio of HC/CC was 1.42 and 1.71-fold increased in *Lcp1* knockout mice at 4 and 8 weeks after ACLT than WT mice (fig.S2, A and B). The area of COL II^+^ (Fig.2, H and I) and ACAN^+^ region (fig.S2, C and D) in *Lcp1* knockout mice were 10.5% to 21.6% higher than in WT mice. The area of MMP13^+^ (Fig.2, J and K), ADAMTS5 ^+^ (fig.S2, E and F) and COL X^+^ region (fig.S2, G and H) in *Lcp1* knockout mice were 3% to 24.5% decreased compared to that of WT mice after ACLT. As osteoclasts mediated sensory nerve infiltration in subchondral bone and pain in OA, we evaluated the effects of *Lcp1* deletion in sensory innervation and pain (*19*). The results showed that the level of NETRIN-1, an essential protein for neural development, increased at 2 weeks after ACLT in WT mice. However, the expression of NETRIN-1 in subchondral area was not observed in *Lcp1* knockout mice (fig.S3, A and B). Correspondingly, the number of CGRP^+^ nerve fibers in *Lcp1*^-/-^ mice were 0.72 and 0.57-fold lower when compared to their littermates (fig.S3, C and D). The von Frey test showed that the threshold of paw withdrawal was 1.7 to 2.6-fold increase in *Lcp1* knockout mice than control mice 3 weeks after ACLT (fig.S3E). To sum up, *Lcp1* knockout alleviates OA progression featured by retarded subchondral bone resorption, alleviated articular cartilage degeneration, and improved pain.

**Fig.2.**
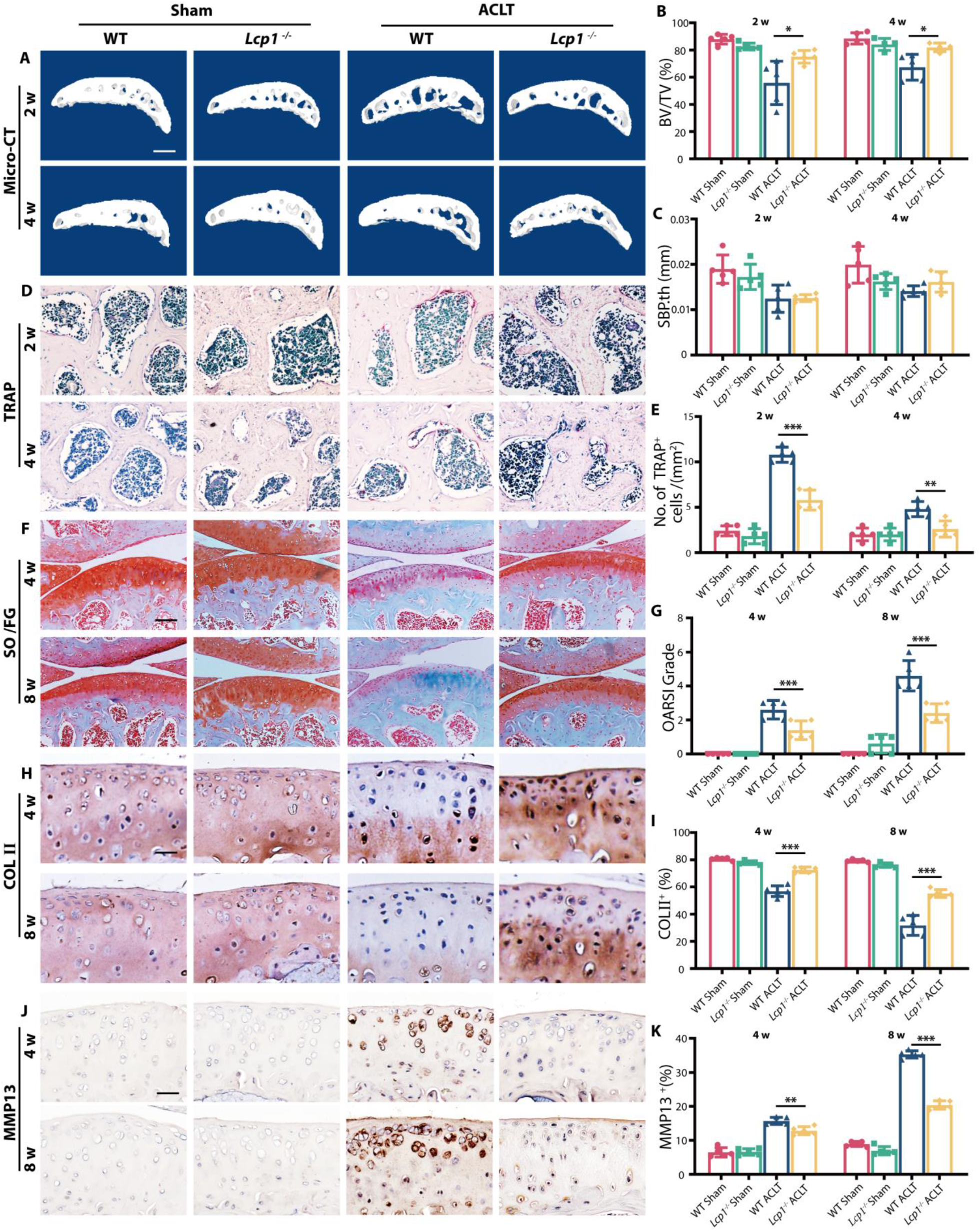
Subchondral bone resorption and articular cartilage degeneration are retarded in Lcp1 knockout mice. **(A)** Representative Micro-CT 3D images of tibia subchondral bone of *Lcp1*^-/-^ mice and WT littermates at 2 and 4 weeks after ACLT. Scale bar, 500μm. **(B-C)** Micro-CT quantitative analysis of tibial subchondral bone, bone volume/tissue volume (BV/TV, %) (B), and subchondral bone plate thickness (SBP. th, mm) (C). **(D)** TRAP staining image of tibial subchondral bone of *Lcp1*^-/-^ mice and WT littermates at 2 and 4 weeks after ACLT. Scale bar,50μm. **(E)** Quantitative analysis of TRAP-positive cells in subchondral bone marrow between *Lcp1*^-/-^ mice and WT littermates. **(F)** Safranin O/Fast Green staining of *Lcp1*^-/-^ mice and WT littermates’ knee articular cartilage. Scale bar, 100μm. **(G)** OARSI grade of knee articular cartilage. **(H)** Representative images of COLII protein immunohistochemistry in tibial articular cartilage of LPL^-/-^ mice and WT littermates at 4 and 8 weeks after ACLT. Scale bar, 20μm. **(I)** Quantitative analysis of COLII protein positive area in articular cartilage. **(J)** Representative images of MMP13 protein immunohistochemistry in tibial articular cartilage of *Lcp1*^-/-^ mice and WT littermates at 4 and 8 weeks after ACLT. Scale bar, 20μm. **(K)** Quantitative analysis of MMP13 protein positive area in articular cartilage. n=5 per group. *P < 0.05, **P < 0.01, and ***P < 0.001.

### *Lcp1* knockout impairs angiogenesis and maintains a low pO_2_ of subchondral bone and cartilage

Under physiological condition, due to lack of blood vessels, the O_2_ concentration in cartilage is strictly maintained at a very low level, for hypoxia was vital for chondrocyte survival and homeostasis (*20*). As osteoclasts mediate type-H vessel formation in subchondral bone after ACLT which provides high oxygen content (*21*). We hypothesized that *Lcp1* knockout inhibited osteoclasts induced angiogenesis and blocked the diffusion of oxygen from subchondral bone to cartilage. To test this hypothesis, we first performed the microangiography of subchondral bone and the results showed that the volume of vessel increased 1.56 and 1.62-fold in WT mice at 4 and 8 weeks after ACLT. In contrast, the vessel volume did not change in *Lcp1*^−/−^ mice after ACLT (0.95 and 1.14-fold compared to sham group) (Fig.3, A and B). Next, we evaluated the level of CD31^hi^EMCN^hi^ vessels. Consistent with the level of vessel volume, the number of CD31 and EMCN positive cells significantly increased to average 20.4 and 23.1 per mm^2^ in WT mice at 4 and 8 weeks after ACLT but increased to average 12.9 and 15.2 per mm^2^ in *Lcp1* deletion after ACLT (Fig.3, C and D). Next, we measured hypoxic status in subchondral bone and cartilage using hypoxia probe. The results of pimonidazole immunostaining, an indicator of hypoxia, revealed that the fluorescence intensity of pimonidazole decreased 33.2% and 81.0% in WT mice at 4 and 8 weeks after ACLT. The fluorescence intensity was maintained in *Lcp1*^−/−^ mice, remained 93.6% and 56.4% intensity of sham group at 4 and 8 weeks (Fig.3, E and F). To further confirm these results, we directly measured the O_2_ levels in the joint in live mice with ^18^F-fluoromisonidazole(^18^F-FMISO)-based PET/CT. Higher ^18^F-FMISO intake indicated lower partial pressure of oxygen (pO_2_) in tissue. The ratio of right (ACLT) and left knee (Sham) uptake of ^18^F-FMISO in WT mice was lower than *Lcp1*^−/−^ mice from 2 weeks after ACLT. At 8 weeks, uptake of ^18^F-FMISO in WT right knee were only 57% to 68% of left knee but ^18^F-FMISO uptake in *Lcp1*^−/−^ mice right knee was 76% to 84% of normal side, indicating the upregulation of pO_2_ in OA progression was retarded after *Lcp1* knockout (Fig.3, G and H). Taken together, *Lcp1* knockout impedes the formation of type-H vessels and maintain the hypoxic environment in subchondral bone and cartilage.

**Fig.3.**
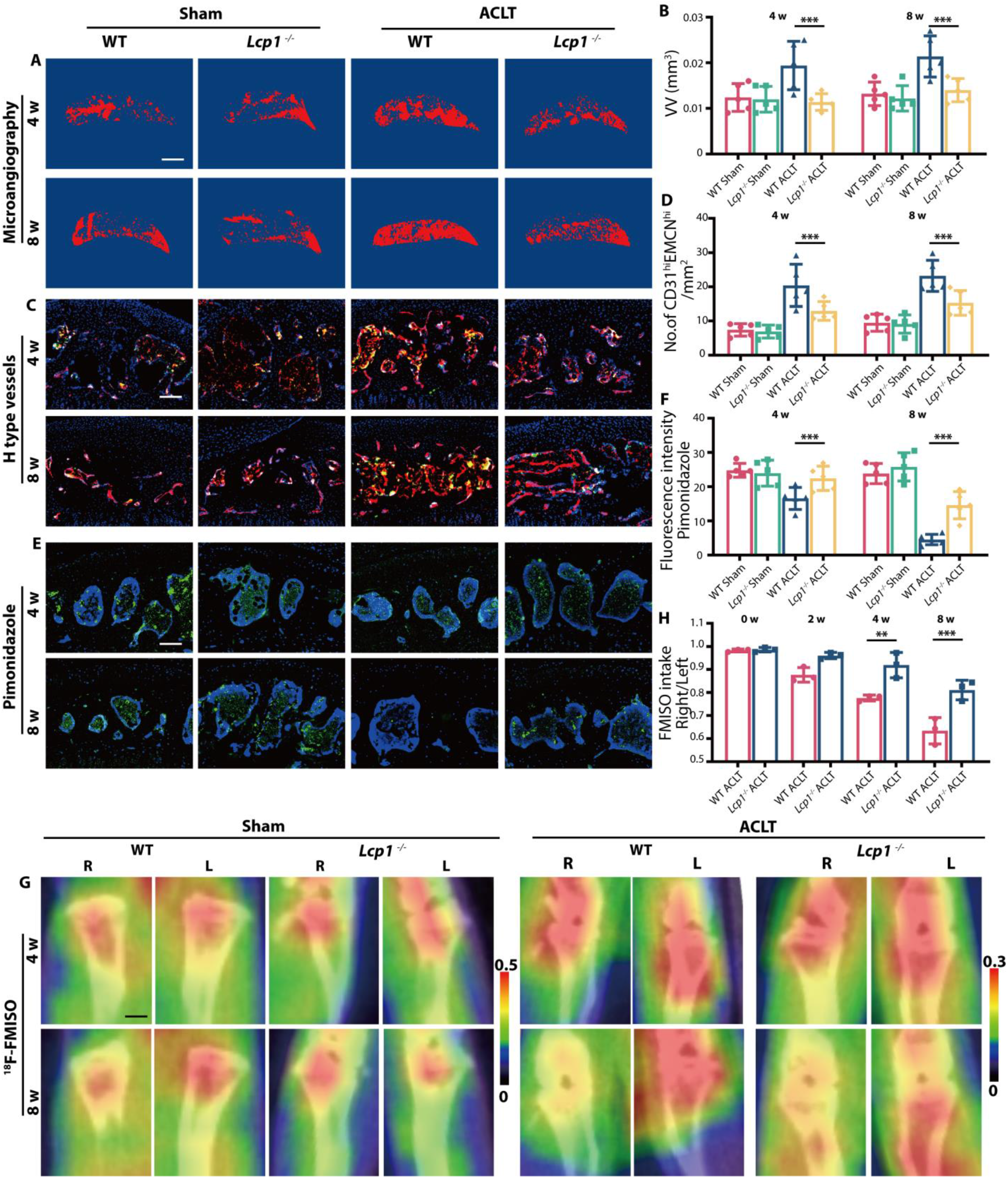
*Lcp1* knockout impairs angiogenesis maintain a low pO2 of subchondral bone and cartilage. **(A)** Three-dimensional image of the sagittal plane of CT-based microangiography in medial tibial subchondral bone of *Lcp1*^-/-^ mice and WT mice at 4 and 8weeks post operation. Scale bar, 500μm. **(B)** Quantification of vessel volume (VV) in medial tibial subchondral bone. **(C)** Maximum intensity projections of immunostaining of endomucin (EMCN) (red), CD31 (green), and Emcn^hi^CD31^hi^ (yellow) cells in medial tibial subchondral bone of *Lcp1*^-/-^ mice and WT mice at 4 and 8weeks post operation. Scale bar, 100μm. **(D)** Quantification of CD31 and Emcn positive cells in subchondral bone marrow. **(E)** Immunostaining of pimonidazole (green) in medial tibial subchondral bone of *Lcp1*^-/-^ mice and WT mice at 4 and 8weeks post operation. Scale bar, 100μm. **(F)** Quantification of pimonidazole fluorescence intensity in subchondral bone marrow. **(G)** ^18^F-FMISO-based PET/CT images of *Lcp1*^-/-^ mice and WT mice at 4 and 8weeks post operation. Scale bar, 10mm. **(H)** Quantification of right knee maximum ^18^F-FMISO uptake/left knee maximum ^18^F-FMISO uptake. n=5 per group. *P < 0.05, **P < 0.01, and ***P < 0.001.

### Hypoxic environment is vital for cartilage maintenance

Next, we explored the mechanism of how pO_2_ affects cartilage degeneration. We hypothesized that the O_2_ affected cartilage chondrocytes through *Hif1-α*, a vital transcriptional factor regulated by oxygen for chondrocytes homeostasis. Tang *et al*. performed single-cell RNA sequencing of human OA cartilage and identified seven chondrocyte populations (*22*). We reanalyzed the database and further divided those cells into six groups (Fig.4A), including fibrocartilage chondrocytes (FCs), OA-associated hypertrophic chondrocytes (OA-HTC), regulatory hypertrophic chondrocytes (rHTC), prehypertrophic chondrocytes (preHTC), homeostatic chondrocytes (HomCs) and proliferative chondrocytes (ProC). We further divided HTC into four clusters (fig.S4A). The results showed that cluster 3 expressed several genes that were responsible for matrix degeneration and endochondral ossification, including *MMP13, COL1A1, COL10A1, RUNX*2, *VEGFC* and *WNT10B* (fig.S4, B-F). Thus, cluster 3 was termed as OA-associated HTC (OA-HTC) which highly expressed *MMP13* and *COL10A1* compared to other subgroups (Fig.4, B and C). GO analysis results showed that OA-HTC participated in replacement ossification, endochondral bone morphogenesis, bone trabecula morphogenesis, embryonic skeletal system development and cartilage development involved in endochondral bone morphogenesis (fig.S5A). KEGG pathway analysis revealed that ECM receptor interaction signaling, and protein digestion and absorption were activated in OA-HTC (fig.S5B). Other clusters of HTCs were defined as regulatory HTC (rHTC) as GO analysis results showed that rHTC regulated calcium channel activity and fatty acid transport (fig.S5A). The GSVA score of different subsets further confirmed that the OA signaling pathway was significantly activated in OA-HTC, more than 1-fold higher than other four groups (Fig.4D). Notably, the HIF1 signaling pathway was downregulated 50% in OA-HTC and FC compared to rHTC, preHTC and ProC, indicating that the downregulation of HIF1 signaling pathway is correlated with OA (Fig.4E).

**Fig.4.**
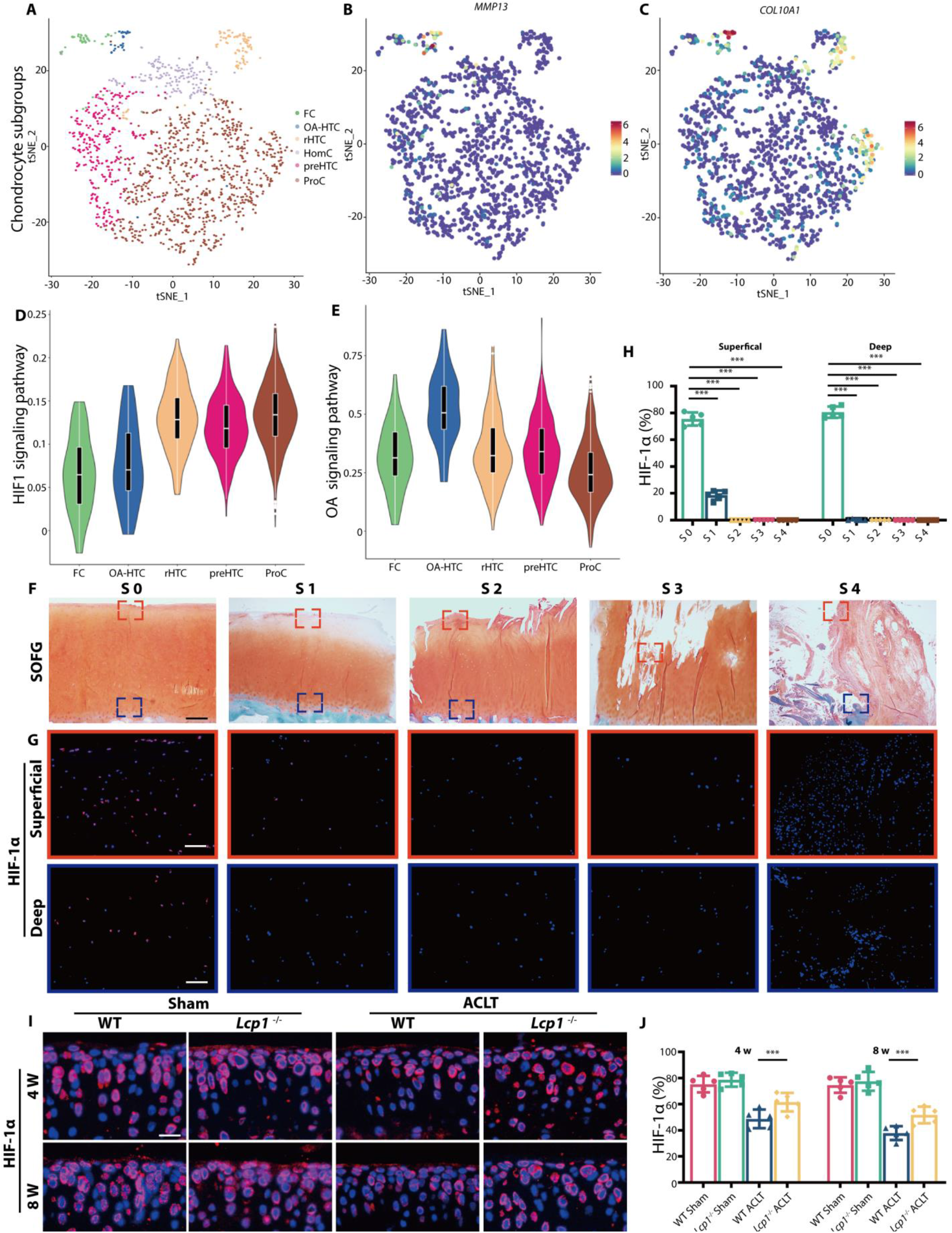
Hypoxic environment is vital for cartilage maintenance. **(A)** Visualization of t-SNE colored according to cell types for human OA cartilage single-cell transcriptomes. **(B-C)** Dot plots showing the expression of *MMP13* and *COL10A1* for OA-associated HTC on the t-SNE map. **(D-E)** Violin plots showing the GSVA score of HIF1 and OA signaling pathway in five major cell types. **(F)** Safranin O/Fast Green staining of different OARSI grade human tibia articular cartilage. Scale bar, 400μm. **(G)** Immunofluorescence staining of HIF-1α protein in different OARSI grade human tibia articular cartilage. Scale bar, 100μm. **(H)** Quantitative analysis of HIF-1α fluorescence intensity in different OARSI grade human articular cartilage. **(I)** Immunofluorescence staining of HIF-1α protein in tibial subchondral bone of *Lcp1*^-/-^ mice and WT littermates at 4 and 8 weeks after ACLT. Scale bar, 20μm. **(J)** Quantitative analysis of HIF-1α fluorescence intensity in *Lcp1*^-/-^ and WT mice articular cartilage. n=5 per group. *P < 0.05, **P < 0.01, and ***P < 0.001.

To further confirm that, we collected human articular cartilage in different OARSI grade from patients (Fig.4F). HIF-1α was abundant in superficial and deep zone in S0 cartilage (80.6% and 75.5%), however, the level substantially decreased when OA progressed (Fig.4, G and H). In S2-S4 phases, HIF-1α expression were hardly detected in cartilage. Similarly, the level of HIF-1α also decreased after ACLT in WT mice and the knockout of *Lcp1* could alleviate this decline. Positive area of HIF-1α in *Lcp1* knockout mice were 1.26 and 1.36-fold higher than WT mice at 4 and 8 weeks after ACLT (Fig.4, I and J). To sum up, the cartilage degeneration is accompanied by the destruction of hypoxia environment and reduced HIF-1α functions and we report a new subtype of HTCs associated with OA.

### Knockdown *Hif 1a* in articular cartilage abolishes protective effect of *Lcp1* knockout

Next, we explored whether *Lcp1* knockout relieved OA progression through inhibiting HIF-1α degradation. We first confirmed that intra-articular injection of adeno-associated virus (AAV) carrying *Hif1a* knockdown shRNA was capable of knocking down *Hif1a* in WT and *Lcp1*^−/−^ mice with 79.2% to 87.6% rate (Fig.5, A and B). After *Hif1a* knocked down, the OARSI grade significantly increased 1.8 and 2.1-fold compared to negative control (NC) in *Lcp1*^−/−^ mice after ACLT (Fig.5, C and D). The ratio of HC/CC decreased 1.1 and 1.2-fold in *Hif1a* AAV mice at 4 and 8 weeks after ACLT compared with NC group (fig.S6, A and B). The area of COL II^+^ (Fig.5, E and F) and ACAN^+^ region (fig.S6, C and D) in *Hif1a* AAV mice substantially decreased 7.9% to 21.3% compared to NC mice, while the area of MMP13^+^(Fig.5, G and H), ADAMTS5 ^+^ (fig.S6, E and F) and COL X^+^ region (fig.S6, G and H) raised 1.44 to 1.93-fold compared to NC mice after operation, indicating destruction of hyaline cartilage greater in *Hif1a* AAV group. Then we checked whether *Hif1a* knockdown in cartilage had effects on subchondral bone. The micro-CT results showed that there was no significant difference of the bone volume or subchondral bone plate thickness between *Hif1a* AAV and negative control mice after ACLT (fig.S7, A to C). Also, there was no statistical difference of the number of TRAP^+^ cells in subchondral bone between two groups (fig.S7, D and E), indicating *Hif1a* knockdown in cartilage did not affect protective effect of subchondral bone in *Lcp1* knockout mice. To sum up, silencing HIF-1α in cartilage abolishes the protective effects of OA progression by *Lcp1* knockout, indicating that inhibiting subchondral bone remodeling alleviates cartilage degeneration through maintaining HIF-1α functions in chondrocytes.

**Fig. 5.**
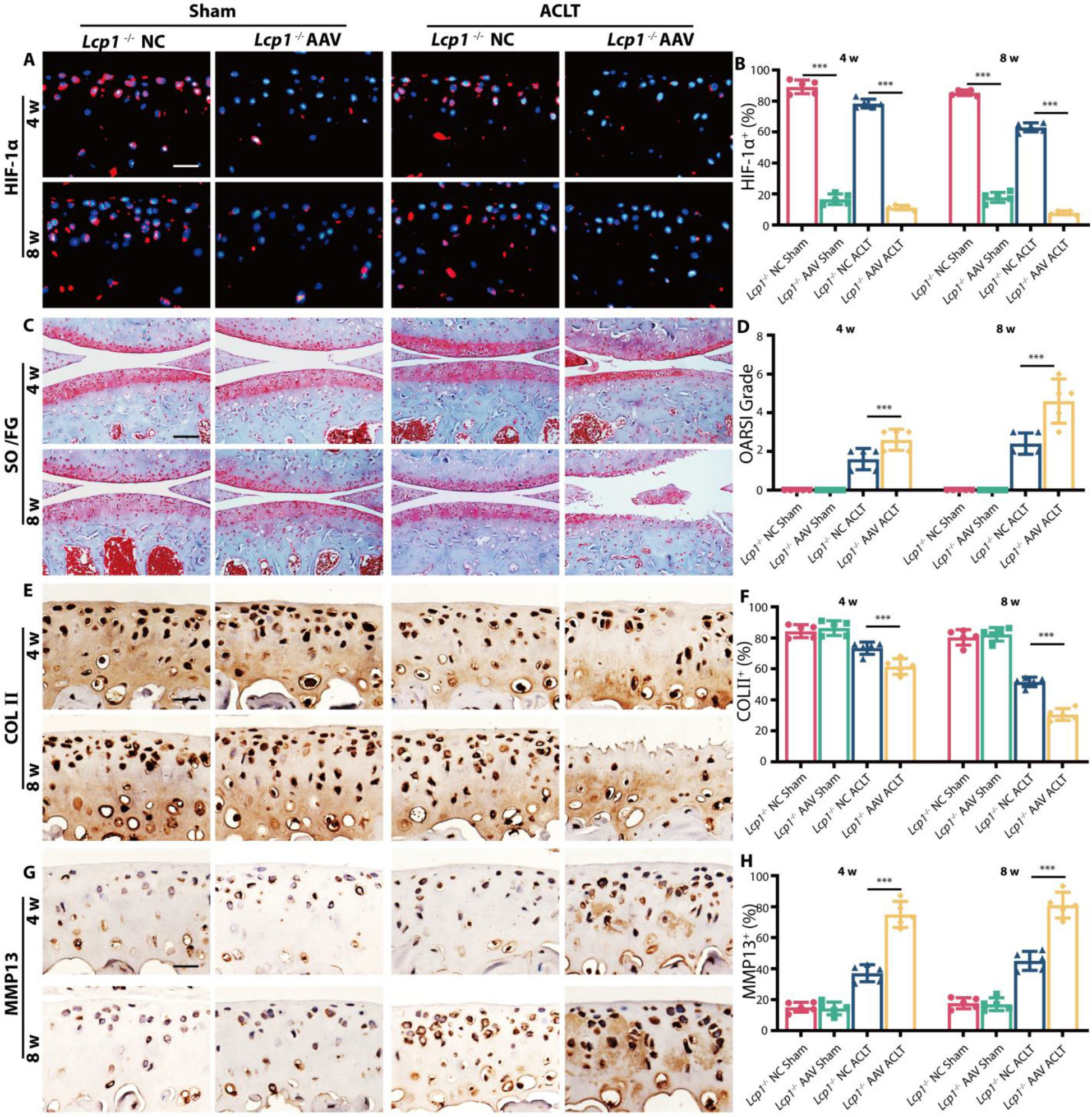
Knockdown *Hif1a* in articular cartilage abolishes protective effect of *Lcp1* knockout. **(A)** Immunofluorescence staining of HIF-1α protein in tibial articular cartilage of ***Lcp1***^-/-^ mice with *Hif1a* AAV or negative control AAV at 4 and 8 weeks after ACLT. Scale bar, 20 μm. **(B)** Quantitative analysis of HIF-1α fluorescence intensity of mice articular cartilage. **(C)** Knee articular cartilage safranin O/Fast Green staining of ***Lcp1***^-/-^ mice with *Hif1a* AAV or negative control AAV at 4 and 8 weeks after ACLT. Scale bar, 100μm. **(D)** OARSI grade of knee articular cartilage. **(E)** Representative images of COLII protein immunohistochemistry in articular cartilage of ***Lcp1***^-/-^ mice with Hif-1α AAV or negative control at 4 and 8 weeks after ACLT. Scale bar =20μm. **(F)** Quantitative analysis of COLII protein positive area in articular cartilage. **(G)** Representative images of MMP13 protein immunohistochemistry in tibial articular cartilage of *Lcp1*^-/-^ mice with *Hif1a* AAV or negative control at 4 and 8 weeks after ACLT. Scale bar, 20μm. **(H)** Quantitative analysis of MMP13 protein positive area in articular cartilage. n=5 per group. *P < 0.05, **P < 0.01, and ***P < 0.001.

### Stabilizing HIF-1α protects articular cartilage in OA

As HIF-1α deficiency worsens OA progression, we speculated that stabilizing HIF-1α could have therapeutic effects on OA. We first confirmed that dimethyloxallyl glycine (DMOG) could preserve 49% to 74% of HIF-1α in ACLT mice at 4 and 8 weeks (Fig.6, A and B). After intraperitoneal injection of DMOG, the OARSI grade significantly decreased 1.6 and 2.1-fold compared to the vehicle group at 4 and 8 weeks after ACLT (Fig.6, C and D). The ratio of HC/CC increased 1.2 and 1.9-fold in DMOG treatment mice at 4 and 8 weeks compared to the vehicle group after ACLT (fig.S8, A and B). Also, the area of COL II^+^ (Fig.6, E and F) and ACAN^+^ region (fig.S8, C and D) in DMOG group substantially increased 2.6% to 18.1% compared to the vehicle group, and the area of MMP13^+^ (Fig.6, G and H), ADAMTS5^+^ (fig.S8, E and F) and COL X^+^ (fig.S8 G and H) decreased 4.7% to 34.7% compared to vehicle mice after operation. Next, we explored whether DMOG had effects on subchondral bone. The results of Micro-CT revealed that there was no significant difference of the bone volume or subchondral bone plate thickness between DMOG and vehicle mice (fig.S9, A to C). Besides, there was no statistical difference of the number of TRAP^+^ cells in subchondral bone between DMOG and vehicle mice with average 7.6 and 8 per mm^2^ at 4 weeks after ACLT (fig.S9, D and E), indicating DMOG had no effects on abnormal bone remodeling in subchondral bone. Above all, stabilizing HIF-1α in chondrocytes could prevent cartilage degeneration in OA regardless of subchondral bone alterations.

**Fig.6.**
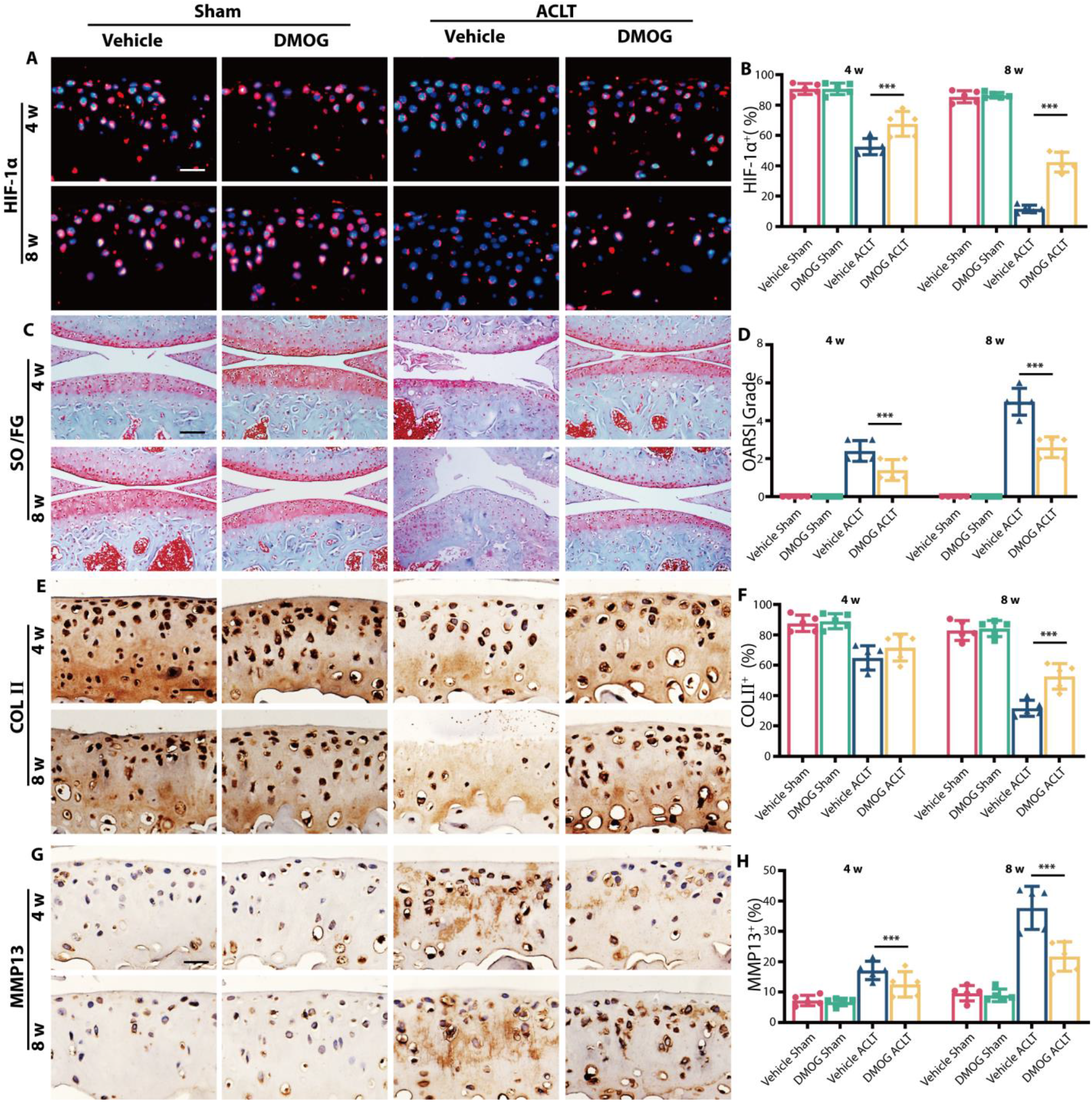
HIF-1α stabilizer DMOG protects articular cartilage in OA. **(A)** Immunofluorescence staining of HIF-1α protein in tibial articular cartilage of WT mice with DMOG or normal saline at 4 and 8 weeks after ACLT. Scale bar, 20μm. **(B)** Quantitative analysis of HIF-1α fluorescence intensity of mice articular cartilage. **(C) K**nee articular cartilage Safranin O/Fast Green staining of WT mice with DMOG or normal saline at 4 and 8 weeks after ACLT. Scale bar, 100μm. **(D)** OARSI grade of knee articular cartilage. **(E)** Representative images of COLII protein immunohistochemistry in articular cartilage of WT mice with DMOG or normal saline at 4 and 8 weeks after ACLT. Scale bar, 20μm. **(F)** Quantitative analysis of COLII protein positive area in articular cartilage. **(G)** Representative images of MMP13 protein immunohistochemistry in tibial articular cartilage of WT mice with DMOG or normal saline at 4 and 8 weeks after ACLT. Scale bar, 20μm. **(H)** Quantitative analysis of MMP13 protein positive area in articular cartilage. n=5 per group. *P < 0.05, **P < 0.01, and ***P < 0.001.

### Oroxylin A alleviates OA progression in WT mice

As *Lcp1* knockout could inhibit OA progression, we then explored whether LPL could serve as a target for OA treatment. Previously, we found that Oroxylin A (Oxy A) was a specific agent targeting LPL (*18*). Therefore, we checked if Oxy A could alleviate OA. After Oxy A treatment, the OARSI grade significantly decreased 1.5 and 2.5-fold compared to the vehicle group at 4 and 8 weeks after ACLT (Fig.7, A and D). The ratio of HC/CC increased 1.3 and 1.7-fold in Oxy A treatment mice at 4 and 8 weeks compared to the vehicle group after ACLT (fig.S10, A and E). Also, the area of COL II^+^ (Fig.6, B and E) and ACAN^+^ region (fig.S10, B and F) in Oxy A group were improved 11.8%-23.1% compared to the vehicle group. The area of MMP13^+^ region (Fig.7, C and F), ADAMTS5^+^ region (Fig.S10, C and G) and COL X^+^ region (Fig.S10, D and H) decreased 4.9% to 37.0% compared to vehicle mice after operation. Next, we explored whether Oxy A had effects on subchondral bone. The results of micro-CT revealed that the bone volume and subchondral bone plate thickness increased at 2 and 4 weeks after ACLT between Oxy A and vehicle mice (fig.S11, A, C and D). The number of TRAP^+^ cells in subchondral bone after Oxy A treatment were 44.8% and 57.1% of vehicle group at 4 and 8 weeks after ACLT (fig.S11 B and E). We evaluated the effects of Oxy A on sensory innervation and pain. The results showed that the level of NETRIN-1 increased at 2 weeks after ACLT in vehicle mice and Oxy A decreased 1.2-fold of the expression of NETRIN-1 at same time (fig.S12, A and C). Consequently, the number of CGRP^+^ nerve fibers decreased 1.3 and 1.4-fold in Oxy A mice compared to the vehicle (fig.S12, B and D). The von Frey test showed that the threshold of paw withdrawal was higher in Oxy A mice than vehicle mice from 4 weeks to 7 weeks (fig.S12E). Above all, targeting LPL is a promising manner to protect subchondral bone and cartilage in OA.

**Fig. 7.**
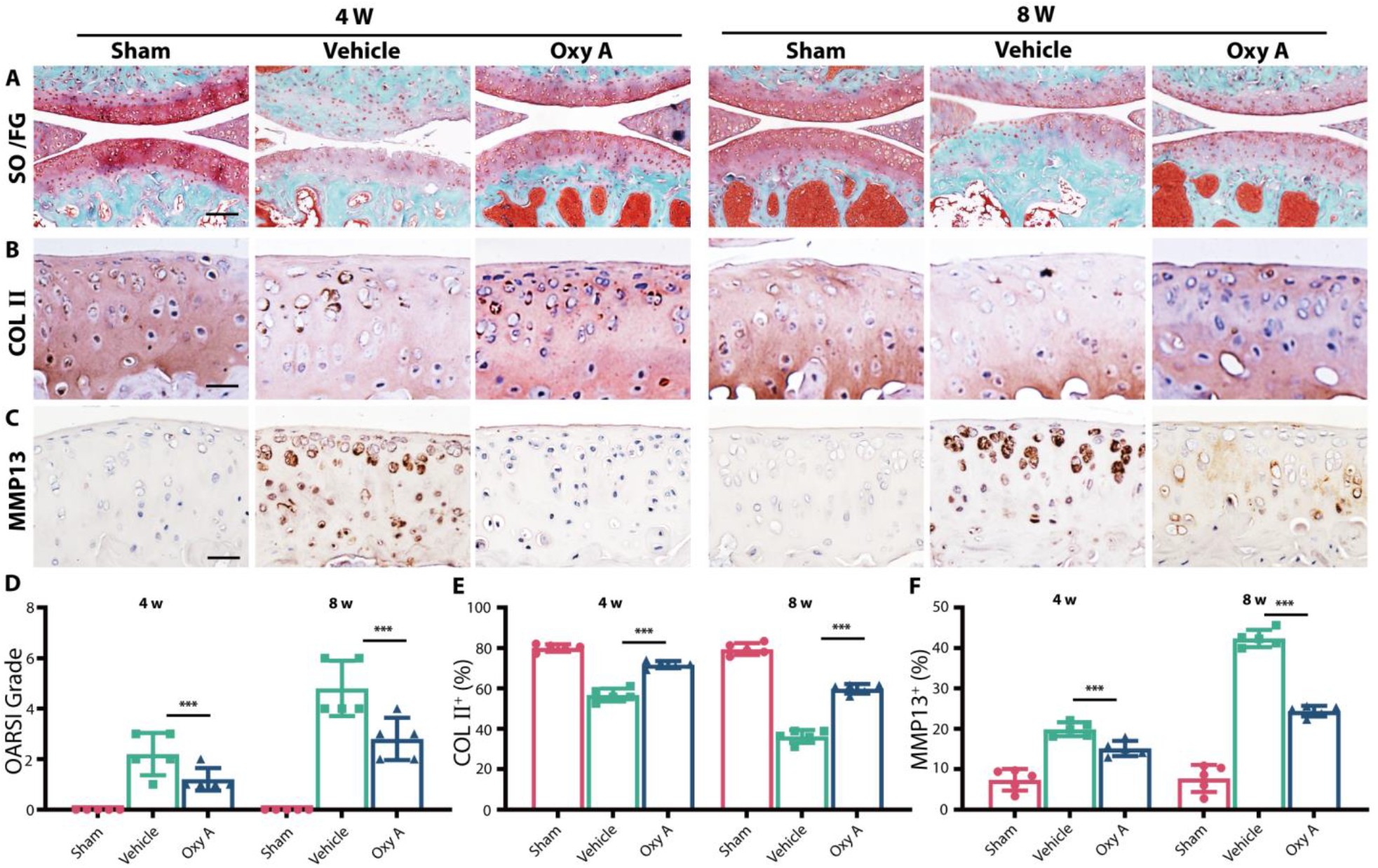
Oroxylin A alleviates OA progression in WT mice. **(A)** Knee articular cartilage Safranin O/Fast Green staining of WT mice with Oxy A or normal saline. Scale bar, 100μm. Representative images of COLII protein immunohistochemistry in tibial articular cartilage of WT mice with Oxy A or normal saline at 4 and 8 weeks after ACLT. Scale bar, 20μm. **(C)** Representative images of MMP13 protein immunohistochemistry in tibial articular cartilage of WT mice with Oxy A or normal saline at 4 and 8 weeks after ACLT. Scale bar, 20μm. **(D)** OARSI grade of knee articular cartilage. **(E)** Quantitative analysis of COLII protein positive area in articular cartilage. **(F)** Quantitative analysis of MMP13 protein positive area in articular cartilage. n=5 per group. *P < 0.05, **P < 0.01, and ***P < 0.001.

**Figure 8.**
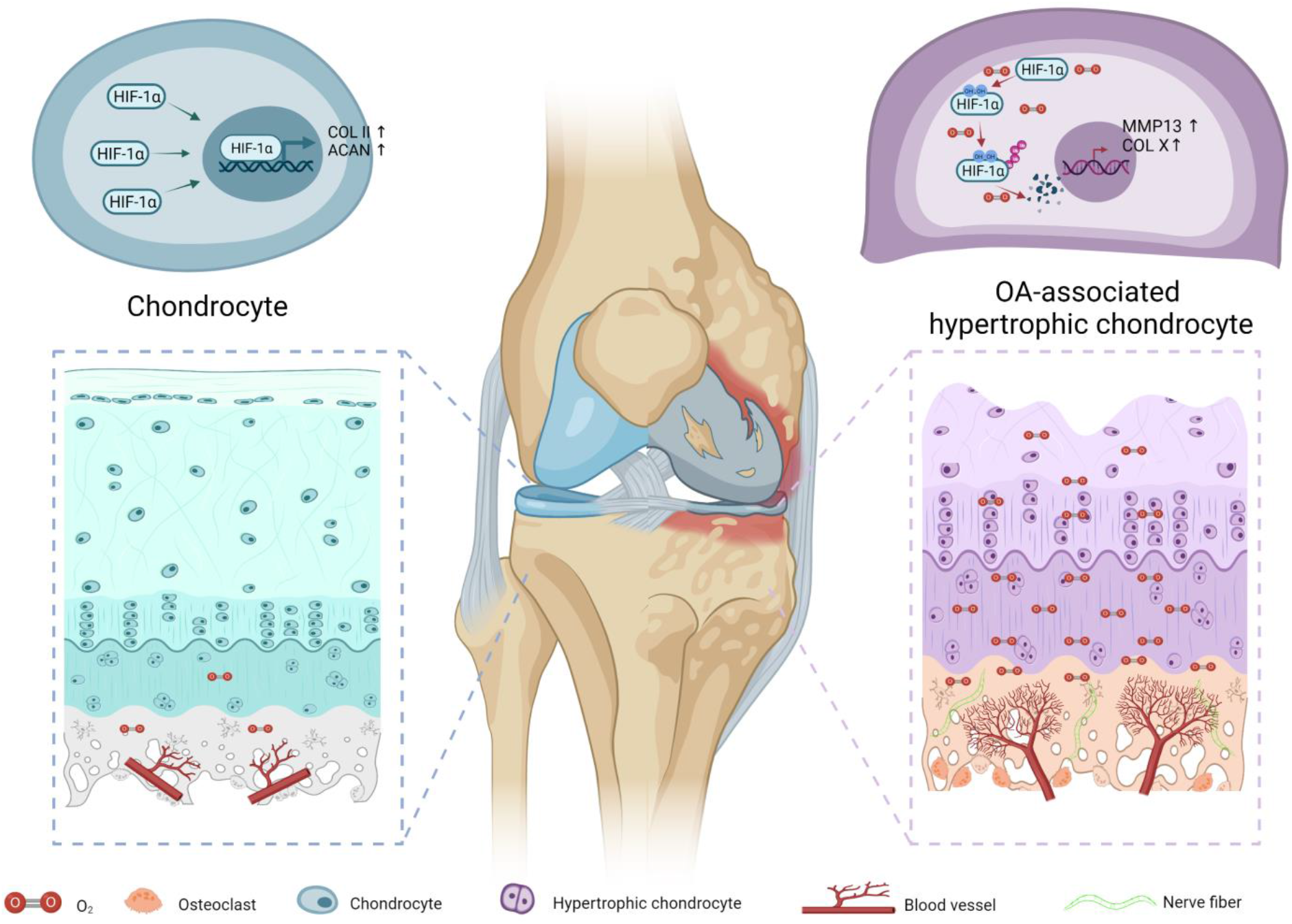
Hypoxic environment is altered by type-H vessel recruited by osteoclasts and high-level oxgen contributes to cartilage degeneration by degrading HIF-1α.

## DISCUSSION

Subchondral bone and cartilage are tightly integrated to form osteochondral units and the relationship between subchondral bone destruction and cartilage degeneration has always been a controversial issue (*23*). Under physiological conditions, subchondral bone maintains a low bone turnover rate and stable microstructure to bear joint load through strictly inhibiting osteoclast formation (*24*). In early OA, abnormal biomechanical and biochemical factors recruit and promote osteoclast differentiation, which results in enhanced bone turnover rate with subchondral bone plate thinning and trabecular bone thickness decreasing (*25*). Conversely, the late OA shows subchondral bone sclerosis characterized by subchondral bone plate and trabecular bone thickening due to excess bone formation (*26*). It is very mysterious and intriguing that the osteoclast shows and disappears, but the cartilage continuously undergoes degeneration. Based on the above, we first hypothesized that osteoclasts introduced certain factors which contribute to sustained cartilage deterioration even without osteoclasts.

Studies showed that targeting osteoclast abnormal activation could block abnormal subchondral bone remodeling in the very beginning and protect articular cartilage degeneration (*27-30*). Previously we reported that LPL, an actin-bundling protein, is indispensable for osteoclast fusion and resorption (*18*). In this study, we first used *Lcp1* knockout mice and performed ACLT to observe whether *Lcp1* knockout could prevent OA progression. Interestingly, although *Lcp1* knockout significantly increased bone volume in bone marrow cavity, it did not affect subchondral bone due to the very low bone turnover rate. To exclude the possible interference of chondrocytes, we showed that LPL was not expressed by chondrocytes. As previously reported, retarding osteoclastogenesis in subchondral bone inhibited the bone turnover rate in early stage of OA and protected the cartilage from degeneration.

How osteoclasts initiate cartilage degeneration is still unclear. In early OA subchondral bone shows increased bone resorption but in advanced OA it shows abnormal bone formation closely correlated to cartilage degeneration believed similar to endochondral ossification (*31, 32*). Dou *et al*. proposed the conception of ‘Osteoclast (OC)-Chondrocyte (CC) crosstalk’ and described several pathways by which these cells might communicate: 1) OC and CC interact via secreted cytokines crossing microsplits and vessels. 2) OC precursors could migrate to the cartilage by invasive vascularization. 3) Mature OC move into subchondral bone and overlying cartilage and interplay with CC in the cartilage area. 4) Subchondral bone deterioration mediated by OC transfers shear forces to the cartilage and subsequently results in aberrant chondrocyte metabolism (*33*). However, no hypothesis could fully explain all. As type-H vessel couples bone resorption and formation (*34*), we considered that subchondral bone type-H vessels could link osteoclasts, osteoblasts, and articular chondrocytes. But insufficient experimental evidence limits further understandings.

Type H vessel, which is rare in normal subchondral bone, is significantly increased in OA induced by osteoclasts (*13, 21, 35*). Type H vessels not only bring secreted mediators and mononuclear cells, but introduce abundant oxygen, an important small molecule in cartilage homeostasis. Normally, the subchondral bone and cartilage are very hypoxic. Level of oxygen is 1∼5% in cartilage and 7% in subchondral bone (*20*). Several invasive tests of subchondral bone local pO_2_ based on mass spectrometry reported various value of pO_2_ from 30-39 mmHg (*36-39*). However, using of oxygen electrode assessment was deficient in spatial resolution, and the implant of the needle electrode may destroy the microvasculature. A direct in vivo measurement of pO_2_ using two-photon phosphorescence lifetime microscopy reported the pO_2_ in bone marrow, but data of subchondral bone was absent (*40*). Thus, we determined the pO_2_ of subchondral bone with a noninvasive and *in vivo* hypoxia probe.

In this study we used ^18^F-FMISO as a hypoxia probe and detected the hypoxia status of subchondral bone and cartilage through PET-CT *in vivo*. ^18^F-FMISO based on the nitroimidazole structure, is the first hypoxia PET tracer used in clinical studies (*41-43*). ^18^F-FMISO is reduced and covalently bound to intracellular macromolecules in hypoxic cells and will not escape from those cells (*44*). Its binding is inversely proportional to the level of oxygen and substantial retention occurs in tissue where oxygen levels below 10 mmHg. We found that in early stage of OA, uptake of ^18^F-FMISO decreased and pressure of O_2_ increased in the ACLT joint. Pimonidazole immunostaining further confirmed the results that pO_2_ increased in early stage of OA. Combining with the results of increased type-H vessels in OA, we believed that type-H vessels induced by osteoclasts altered the hypoxia environment of subchondral bone and cartilage.

Then we explored how increased oxygen affected chondrocytes. HIF-1α is of critical significance in cartilage energy metabolism, differentiation fate and matrix homeostasis. HIF-1α serves as a pivotal factor for chondrocytes by strengthening anaerobic glycolysis and impeding apoptosis via mitophagy (*45, 46*). As a vital chondrocyte marker, *SOX9* is regulated by HIF-1α and could inhibit hypertrophic alterations of chondrocytes (*15, 47, 48*). Hypoxia and HIF-1α Induce extracellular matrix synthesis through promoting the expression of *COL2A1*, and *ACAN*, and inhibiting the expression of *COL1A1, COL1A2, COL10A1* and *MMP13*, which renew articular cartilage matrix in a harmonious and stable rate (*49-52*).

To further testify the roles of HIF-1α in OA, we reanalyzed the single cell RNA-sequencing data of OA human chondrocytes from Tang (GSE104782) (*22*). Tang obtained cartilage samples from knee joints of 16 patients with trauma and 16 with rheumatoid arthritis. They empirically defined 4 populations of chondrocytes: ProCs, preHTCs, HTCs, FCs and three novel populations, effector chondrocytes (ECs), regulatory chondrocytes (RegCs) and HomCs. We focused on HTCs, a group of cells involved in endochondral ossification and calcified cartilage duplication in OA progression. We found that HTCs had two different subgroups. A subgroup of HTCs which expressed high levels of *MMP13, RUNX2, COL1A1* and *COL10A1* was named OA-associated HTC. KEGG and GO enrichment analysis indicated that OA-associated HTC were involved in endochondral ossification, ossification replacement and endochondral bone morphogenesis, while another subgroup of HTCs was named regulatory HTC involved in regulation of calcium channel activity, AMPA receptor activity and circadian sleep wake cycle. Next, we used GSVA analysis to compare the signal pathway activation in different cell types. OA-associated HTC showed highest score in OA signaling pathway and the second lowest score in HIF-1 pathway. Immunofluorescence of HIF-1α in human knee articular cartilage at different stages of OA and PTOA mice showed that HIF-1α in deep layer was degraded.

We used *Hif1a* knockdown AAV and HIF-1α stabilizer DMOG in *Lcp1* knockout and WT OA mice to verify that HIF-1α can influence articular cartilage independently from subchondral bone alteration. Though *Lcp1* knockout reduced the bone remodeling in subchondral bone, silencing HIF-1α mediated by *Hif1a* AAV could still cause articular cartilage degeneration and abolish the protective effect in *Lcp1*^*-/-*^ ACLT mice. On the other hand, although subchondral bone remodeling was not inhibited in WT ACLT mice, but DMOG stabilized HIF-1α and retarded OA progression in a non-hypoxic environment, consistent with a previous study by Hu *et*.*al* (*46*). HIF-1α could be a potential target in middle or late stages of OA and has wider treatment timing than drugs which target bone remodeling only.

Oxy A could target LPL (*18*). We showed that intraperitoneal injection of Oxy A could inhibit osteoclast formation in OA subchondral bone, reduce cartilage degradation, inhibit CGRP^+^ sensory nerve fiber invasion and relieve joint pain, indicating that Oxy A could be a DMOAD.

Some limitations of this study should be addressed. First, we unveiled the role of HIF-1α stabilizer in OA treatment in rodents, however, its therapeutic effect in human should be further explored. Besides, systemic administration of HIF-1α stabilizer may lead to off-target effect. Local injection with suitable drug delivery system would further improve the outcomes. The last, Oxy A is hydrophobic small molecule which decrease the bioavailability via systematic administration. Several biological materials including hydrogels, nanozyme and nanoparticle has been reported as promising carriers in OA treatment (*53-56*). Thus, we have developed a long stranded, cartilage targeted and enzyme responded biological materials contain a DMOAG to improve the therapeutic effects on cartilage protection in the further study.

### The highlights of this study are as follow

1. In subchondral bone, type H vessels induced by osteoclasts in early OA elevate pO_2_ levels.
2. Increased pO_2_ levels abolish HIF-1α functions of maintaining chondrocytes homeostasis.
3. A new subgroup of HTCs with decreased HIF-1α activity is highly associated with OA progression.

## MATERIALS AND METHODS

### Study design

This study was performed to explore the roles and mechanisms of osteoclasts in subchondral bone area in articular chondrocyte degeneration in OA. First, we used *Lcp1* knockout mice with inhibited osteoclastogenesis in subchondral bone to establish ACLT OA model. We compared the difference of cartilage degeneration, subchondral bone remodeling, angiogenesis and hypoxia environment change in subchondral bone between *Lcp1* knockout mice and WT mice in ACLT model through immunofluorescence and histomorphometric analyses. ^18^F-FMISO PET/CT analysis was used to detect hypoxic environment of knee joint *in vivo* at different time after ACLT. The vital role of HIF-1α in maintaining chondrocyte stability and preventing hypertrophy was verified by single-cell RNA-sequencing. *Hif1a* knock down AAV and HIF-1α stabilizer were used to confirm that *Lcp1* knockout protected cartilage through stabilizing HIF-1α in chondrocytes. Potentiality of LPL as a therapeutic target of OA was testified by systematic administration of Oxy A, which was proved as an LPL specific inhibitor in our former study. Samples were randomly assigned into distinct intervention groups and littermates were included in the control group. Five samples were used for statistical analysis in each experiment. The study was approved by Shanghai Model Organisms (SCXK [Shanghai] 2017-0010 and SYXK [Shanghai] 2017-0012) and IACUC guidelines were followed for animal experiments.

### Mouse models

*Lcp1* knockout mice on C57BL/6 background were created by the Shanghai Model Organisms in our former study (*18*). Male C57 mice (8-week-old) were got from Weitonglihua Corporation (Beijing, China). The rodent researches were carried out in the pathogen-free environment.

Laboratory conditions for mice were listed below. Temperature: 22°C; humidity: 50%; light-dark cycle: 12h; water and food: available. In line with our previous protocol, ACLT surgery was performed to generate OA mouse model. Briefly, after anesthesia with pentobarbital sodium, a longitudinal cutaneous incision was made at medial side of the right knee. The ACL was transected after open knee joint through medial approach of ligamentum patellae under a surgical microscope. The rodents were randomly divided into different groups: Sham (performed the incision without ACL transection), ACLT, group of different intervention (ACLT mice intraarticularly injected with *Hif1a* AAV and DMOG or Oxy A intraperitoneally), and vehicle (ACLT mice injected with saline).

### Human samples

Human samples of medial tibia plateau were collected during total knee arthroplasty operations. Subchondral bone and articular cartilage samples were cut into 1–2 cm pieces and fixed in 4% PFA solution (G1101-500ML Servicebio, Wuhan, China) for 2 days and decalcify in 10% EDTA (G1105-500ML, Servicebio, Wuhan, China) for 6 mouths. Samples were embedded in paraffins or optimal cutting temperature compound (OCT) and cut into 5 μm sections. The experiments were approved by Shanghai University (ECSHU 2021-146).

### *Hif1a* knockdown AAV construction

Recombinant AAV particles were produced using the AAV Helper-Free System (*57*). Step 1: The cloning of the exogenous gene inserted into a suitable vector. In most cases, the exogenous gene was cloned into a vector containing the ITR/MCS vector. The inverted terminal repeat (ITR) sequences in these vectors provided all the cis-acting elements necessary for AAV replication and packaging. Step 2: The recombinant expression plasmids were co-transfected with pHelper (bearing the adenovirus-derived gene) and pAAV-RC (maintaining the AAV replication and capsid genes) into AAV-293 cells (providing the trans-acting elements required for AAV replication and packaging). After 2 to 3 days of transfection, recombinant AAV was assembled in packaging cells. Step 3: AAV particles were collected from infected AAV-293 cells. Step 4: Concentrate and purify the virus from step 3. Step 5: The titer of the resulting virus is determined by quantitative PCR, which gives the physical titer of the AAV genome packaged into the pellet. *Hif1a* knockdown shRNA sequence:5′-GUGGAUAGCGAUAUGGUCAUU-3′(*58*).

### ^18^F-FMISO PETCT analyses

^18^F-FMISO construction: ^18^F-FMISO was synthetized as stated by the means proposed by Yu and He (*44, 59*). The 1-(2-nitro-1′-imidazolyl)-2-O-tetrahydropyranyl-3-O-toluenesulfonylpropanediol precursor was obtained from ABX GmbH (Radeberg, Germany), and ^18^fluoride was got from BV Cyclotron VU (Amsterdam, Netherlands). Radio synthesis was completed in an automated synthesizer. Male mice of the same age were intravenous injected with 100 MBq ^18^F-FMISO in 0.2 ml normal saline. Mice were detected in PET/CT 1 h after the injection of ^18^F-FMISO. Images were collected and analyzed by Bee DICOM Viewer.

### Micro-CT analyses

Tibia subchondral bone vasculature was evaluated by Micro-CT as reported earlier (*30*). Mice were anesthetized with pentobarbital sodium, tibia subchondral bone vessels were washed with normal saline solution, 4% PFA solution and normal saline solution through heart in proper sequence. Then the intravascular contrast media (MICROFIL, MV-120, Flow Tech) was injected. The mice were preserved at 4°C for 12 h before the knee joints were harvested. The knee joints were fixed for 3 days in 4% PFA and decalcified in 10% EDTA for 21 days prior to scan. The vascular volume in subchondral bone was analyzed. The region of interest was central of medial tibia plateau.

### Microangiography

Tibia subchondral bone vasculature was evaluated by Micro-CT as reported earlier (*30*). Mice were anesthetized with pentobarbital sodium, tibia subchondral bone vessels were washed with normal saline solution, 4% PFA solution and normal saline solution through heart in proper sequence. Then the intravascular contrast media (MICROFIL, MV-120, Flow Tech) was injected. The mice were preserved at 4°C for 12 h before the knee joints were harvested. The knee joints were fixed for 3 days in 4% PFA and decalcified in 10% EDTA for 21 days prior to scan. The vascular volume in subchondral bone was analyzed. The region of interest was central of medial tibia plateau.

### Histological analysis

Knee joints were collected and fixed in 4% PFA for 2 d, and decalcified in 10% EDTA for 14 d. Next, the joints were embedded in paraffin or OCT and serially sectioned 5-μm in the sagittal plane of in central the medial compartment of the joints. Then, hematoxylin and eosin (H&E, G1005-500ML, Servicebio, Wuhan, China), Safranin O and fast green (G1053-100ML Servicebio, Wuhan, China), and TRAP staining (G1050-50T, Servicebio, Wuhan, China) were done according to regular procedures *(18,30)*. A light microscope (Olympus BX53) was used for imaging. The tidemark line labeled the bound between HC and CC. H&E staining image was employed to evaluate the thickness of HC and CC. The OARSI grade was used to analyze the degradation of tibial plateau cartilage. The TRAP staining was used to count osteoclasts in subchondral bone (*60*).

### Immunofluorescence and histomorphometry

Antibodies against ACAN (1:500; Servicebio, GB11373) and COLII (1:500; Servicebio, GB11021) were obtained from Servicebio (Wuhan, China). MMP13 (1:200; Proteintech, 18165-1-AP) was purchased from Proteintech (Wuhan, China). LPL (1:200; Thermo, PA5-85216) and ADAMTS5 (1:500; Thermo, PA5-27165) were obtained from Thermo (Waltham, USA). COLX (1:500; Abcam, ab260040), CD31 (1:200; Abcam, ab182981) and CGRP (1:200; Abcam, ab36001) were obtained from Abcam (Cambridge, UK). EMCN (1:50; Santa Cruz, sc-65495) was obtained from Santa Cruz (Dallas, USA). HIF-1α (1:50; Novus, NB100-105) and NETRIN-1 (1:100; Novus, NB100-1605) were purchased from Novus (Littleton, USA). Paraffins sections were dewaxed in dimethylbenzene for 10 min twice and dehydration was performed in 100%, 100%, 95%, 80% and 75% ethanol for 5 min each time. Antigen retrieval was performed in EDTA (9.0 pH) or citrate (6.0 pH) at 95°C for 20 min. Until sections cooled down to room temperature, 3% H_2_O_2_ was used to block peroxidase for 15 min. 10% goat serum was employed to block sections for 1 h. Primary antibody were diluted into appropriate concentration according to instructions and incubated for 12 h at 4°C. After primary antibody was washed three times, HRP conjugated secondary antibody or fluorescein conjugated secondary antibody was incubated for 30-60 min at room temperature according to the manual. DAB kit and hematoxylin were used in immunochemistry staining. DAPI was used to stain nucleus in immunofluorescence. A light microscope (Olympus BX53) was used for section imaging. The number of positive cells or area in sections were quantified using ImageJ.

### Hypoxia probe detection

Hypoxyprobe green kit (hpi, HP6-100) was purchased from Hypoxyprobe. Pimonidazole 60 mg/kg was intraperitoneal injected into mice 1 hour before sacrifice. Paraffin sections were prepared as described above. Primary antibody conjugated with FITC in kit was used to detect pimonidazole at 1:100 dilution.

### Single-cell RNA-seq data analyses

We downloaded Single-cell RNA-seq data from NCBI (GSE10478) and processed the unique molecular identifier (UMI) count matrix with the R package Seurat (version 3.1.1) (*61*). 1600 cells were obtained for following evaluation. Library size normalization was done using Normalize Data function in Seurat to acquire normalized number.

Top variable genes were verified by the means reported by Macosko *et al*. (*62*). The most variable genes were collected using Find Variable Genes function in Seurat. Principal component analysis (PCA) was done to depress the dimensionality with RunPCA function in Seurat (*61*). To cluster cells, graph-based clustering was done based on each cell gene expression profile via the Find Clusters function. Visualization of cells were done by a 2-dimensional t-distributed stochastic neighbor embedding (t-SNE) algorithm using the RunTSNE function in Seurat. Find All Markers function were used to verify marker genes of each cluster. Find All Markers determined positive markers compared to other cells.

Differentially expressed genes (DEGs) were established via the Find Markers function in Seurat (*58*). Significant DEGs threshold were P value < 0.05 and |log2 fold change| > 0.5. GO enrichment and KEGG pathway enrichment analysis of DEGs were respectively carried out by R based on the hypergeometric distribution.

### Von Frey test

Von Frey filaments (Touch test, USA) were used to measure the 50% paw withdrawal threshold (50% PWT) as our previous study. Briefly, mice were put into cages for 30 min to adapt. Von Frey hairs with different forces (gram=0.04, 0.07, 0.16, 0.4, 0.6, 1.0, 1.4, 2.0) were used in this test, and 2.0 g was served as the cutoff threshold. Mechanical allodynia was analyzed based the up-down theory from Dixon and was performed each week (*63*). The filament was stabbed under the middle plantar area of the right hind paw. Negative response was record as “o” and higher force was used in next test. Positive response was record as “x”, and next lower force was employed. When the difference of response occurred (“ox” or “xo”), four more experiments were done to get six results. The interval between neighboring stab was 6 min. 50% PWT was analyzed by the formula: 10[Xf + kδ] /10^4^, where Xf is the value of the last force applied is a constant of serial force and k is derived from response pattern.

### Statistical analyses

All data were presented as the mean ± SD or SEM. Comparisons between two groups were performed by the two-tailed Student’s t test. Comparisons among multiple groups were analyzed by the one-way analysis of variance (ANOVA). The results were visualized and analyzed by the GraphPad PRISM software, and a P < 0.05 implied differences were statistically significant.

## Supplementary Materials

**Fig.S1 L-plastin positive cells distribution in proximal tibia of sham and ACLT mice**

**Fig.S2 Calcified cartilage duplication is retarded in Lcp1 knockout mice**.

**Fig.S3 Lcp1 knockout mice show decreased NETRIN-1 and CGRP+ sensory nerves in the subchondral bone and pain amelioration**.

**Fig.S4 HTC subsets definition and characteristic of OA-associated HTC**.

**Fig.S5 KEGG and GO analysis of ProC, PreHTC, rHTC, OA-HTC and FC**.

**Fig.S6 Knockdown Hif-1a abolishes cartilage protective effects in Lcp1-/-mice**.

**Fig.S7 Knockdown Hif-1a in Lcp1-/-mice shows no effects on subchondral bone**.

**Fig.S8. DMOG prevents calcified cartilage duplication**.

**Fig.S9 DMOG does not affect subchondral bone in OA**.

**Fig.S10. Calcified cartilage duplication retarded after Oroxylin A injection**.

**Fig.S11 Oroxylin A inhibits subchondral bone remodeling via inhibiting osteoclast formation**.

**Fig.S12. Oroxylin A decreases Netrin-1 and CGRP+ sensory nerves in the subchondral bone and ameliorates pain in WT mice**.

**Movies S1 Video of** ^**18**^**F-FMISO PETCT in *Lcp*1**^**-/-**^ **mice 4 weeks after ACLT**

**Movies S2 Video of** ^**18**^**F-FMISO PETCT in WT mice 4 weeks after ACLT**

## Acknowledgments

We thank the Shanghai Model Organisms for constructing *Lcp1* knockout mice. We thank department of nuclear medicine of Renji hospital affiliated to Shanghai JiaoTong University school of Medicine for providing ^18^F-FMISO and PET/CT. We thank OBiO Technology (Shanghai) for constructing *Hif1a* knockdown AAV. We thank OE biotech company (Shanghai, China) for the supporting of bioinformatics analysis.

## Author contributions

Conceptualization: H.Z, LP.W, J.C,X.C,. Methodology: H.Z, HD.S, C.W, J.C. Investigation: H.Z, SC.W, QR.Z, JW.G, XC.Z, SH.S, T.Z. Visualization: H.Z, SC.W, DY.Z, YF.H, J.C, FX.W, QM,G. Funding acquisition: YY.J, X.C, JC.S. Project administration: YY.J, X.C. Supervision: YY.J, X.C, JC.S. Writing – original draft: H.Z, LP.W. Writing – review & editing: YY.J, X.C, JC.S.

## Data and materials availability

All data associated with this study are present in the paper and/or the Supplementary Materials. Additional data related to this paper may be requested from the authors.

