## supplemental material for "Maintaining hypoxia environment of subchondral bone alleviates osteoarthritis progression"

**This PDF file includes:**

Fig.S1 L-plastin positive cells distribution in proximal tibia of sham and ACLT mice
Fig.S2 Calcified cartilage duplication is retarded in Lcp1 knockout mice.
Fig.S3 *Lcp1* knockout mice show decreased NETRIN-1 and CGRP+ sensory nerves in the subchondral bone and pain amelioration.
Fig.S4 HTC subsets definition and characteristic of OA-associated HTC.
Fig.S5 KEGG and GO analysis of ProC, PreHTC, rHTC, OA-HTC and FC.
Fig.S6 Knockdown Hif-1a abolishes cartilage protective effects in *Lcp1*$^{-/-}$ mice.
Fig.S7 Knockdown Hif-1a in *Lcp1*$^{-/-}$ mice shows no effects on subchondral bone.
Fig.S8. DMOG prevents calcified cartilage duplication.
Fig.S9 DMOG does not affect subchondral bone in OA.
Fig.S10. Calcified cartilage duplication retarded after Oroxylin A injection.
Fig.S11 Oroxylin A inhibits subchondral bone remodeling via inhibiting osteoclast formation.
Fig.S12. Oroxylin A decreases Netrin-1 and CGRP+ sensory nerves in the subchondral bone and ameliorates pain in WT mice.

**Other Supplementary Materials for this manuscript include the following:**

[use this section only if you have movies, audio or data files]

Movies S1 Video of $^{18}F$-FMISO PETCT in *Lcp1*$^{-/-}$ mice 4 weeks after ACLT

Movies S2 Video of $^{18}F$-FMISO PETCT in WT mice 4 weeks after ACLT

### MATERIALS AND METHODS

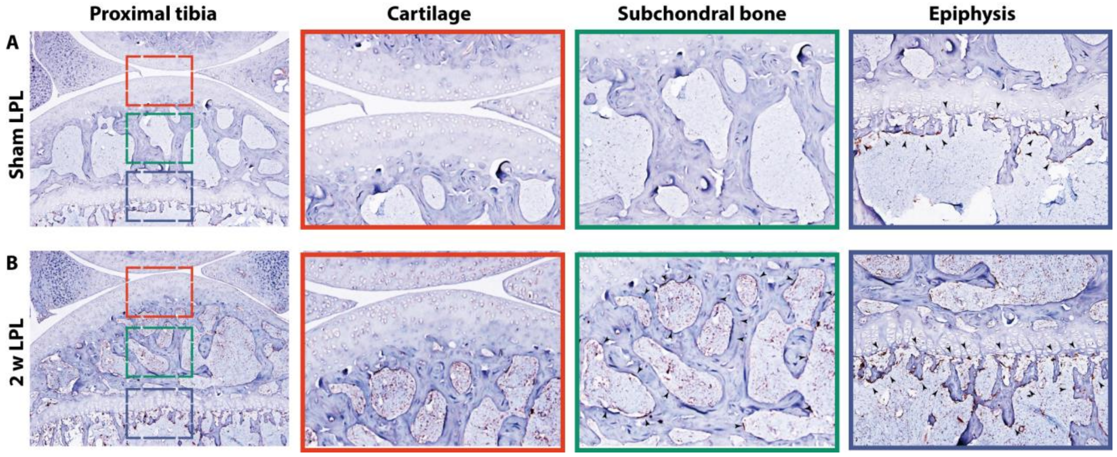


**Fig.S1 L-plastin positive cells distribution in proximal tibia of sham and ACLT mice**

**(A)** Representative images of LPL protein immunohistochemistry in whole normal proximal tibia, articular cartilage, subchondral bone, and epiphysis. **(B)** Representative images of LPL protein immunohistochemistry in whole proximal tibia, articular cartilage, subchondral bone, and epiphysis 2 weeks after ACLT. Scale bar, 100 μm (first column). Scale bar, 50 μm (2-4 column).

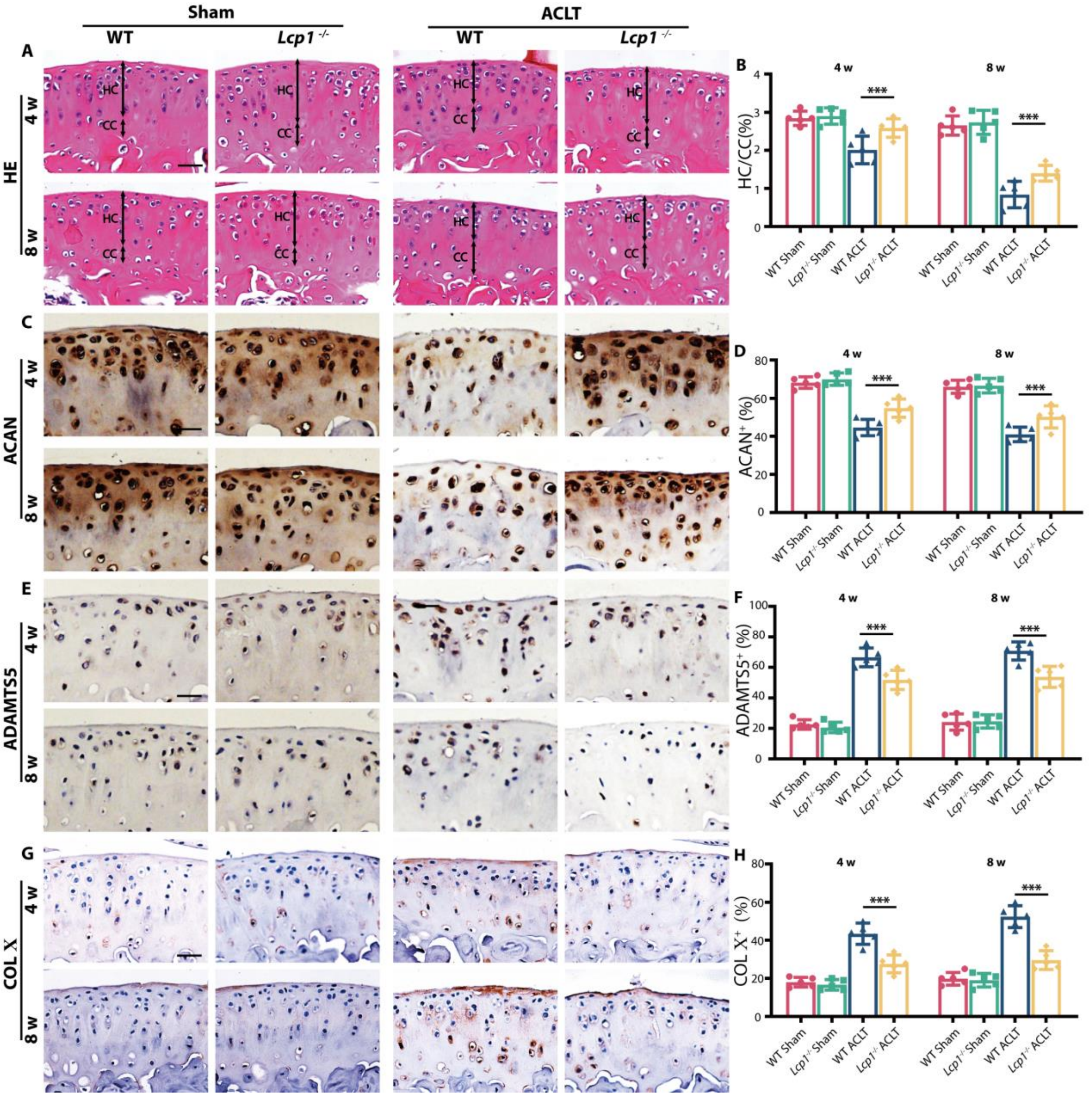


**Fig.S2 Calcified cartilage duplication is retarded in *Lcp1* knockout mice.**

**(A)** Representative images of Hematoxylin and eosin staining of proximal tibia articular cartilage of *Lcp1*$^{-/-}$ mice and WT littermates at 4 and 8 weeks after ACLT. Double-headed arrows label range of HC and CC. Scale bar, 50μm. **(B)** Quantitative analysis of HC and CC thickness ratio. **(C)** Representative images of ACAN protein immunohistochemistry in tibial articular cartilage of *Lcp1*$^{-/-}$ mice and WT littermates at 4 and 8 weeks after ACLT. Scale bar, 20μm. **(D)** Quantitative analysis of ACAN protein positive area in articular cartilage. **(E)** Representative images of ADAMTS5 protein immunohistochemistry in tibial articular cartilage of *Lcp1*$^{-/-}$ mice and WT littermates at 4 and 8 weeks after ACLT. Scale bar =20μm. **(F)** Quantitative analysis of ADAMTS5 protein positive area in articular cartilage. **(G)** Representative images of COL X protein immunohistochemistry in tibial articular cartilage of *Lcp1*$^{-/-}$ mice and WT littermates at 4 and 8 weeks after ACLT. Scale bar =20μm. **(H)** Quantitative analysis of COL X protein positive area in articular cartilage. N=5 per group. *P < 0.05, **P < 0.01 and ***P < 0.001.

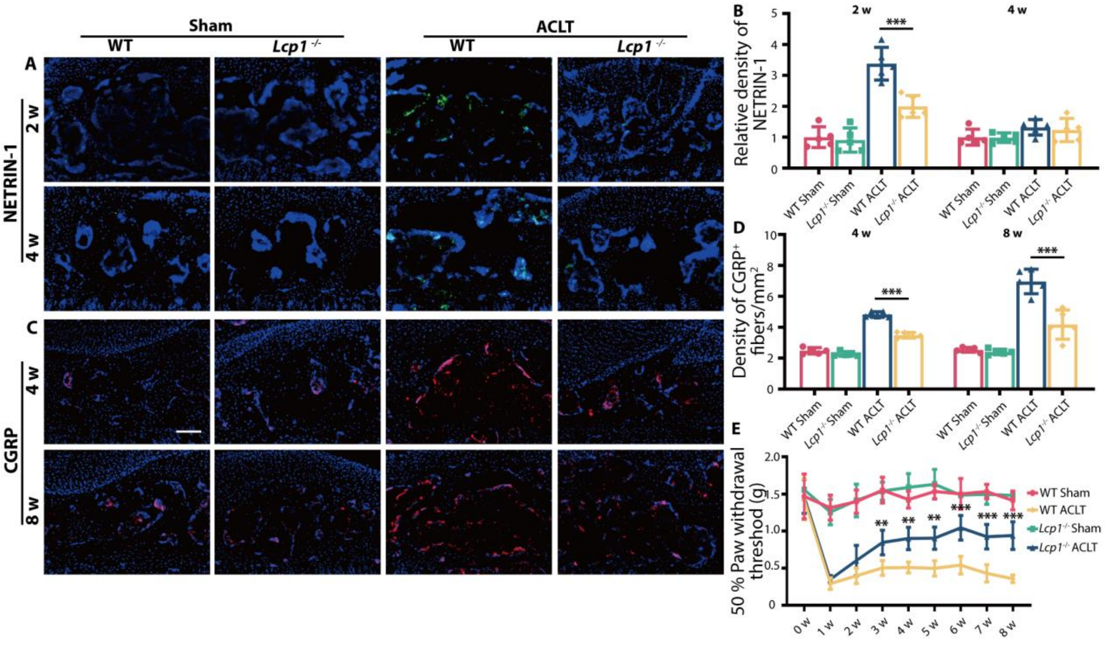


**Fig.S3 *Lcp1* knockout mice show decreased NETRIN-1 and CGRP$^{+}$ sensory nerves in the subchondral bone and pain amelioration.**

**(A)** Immunofluorescence staining of NETRIN-1 protein in *Lcp1*$^{-/-}$ and WT mouse tibial subchondral bone 2 and 4 weeks after ACLT. Scale bars, 100 µm. **(B)** Quantitative analysis of density of NETRIN-1 in subchondral bone marrow. **(C)** Immunofluorescence staining of CGRP$^{+}$ sensory nerve fibers in *Lcp1*$^{-/-}$ and WT mouse tibial subchondral bone 4 and 8 weeks after ACLT surgery. Scale bars, 100 µm. **(D)** Quantitative analysis of the density CGRP$^{+}$ nerve fibers in subchondral bone marrow. **(E)** Paw withdrawal threshold was tested at the right hind paw of *Lcp1*$^{-/-}$ and WT each week after surgery until 8 weeks. N=5 per group. *P < 0.05, **P < 0.01 and ***P < 0.001.

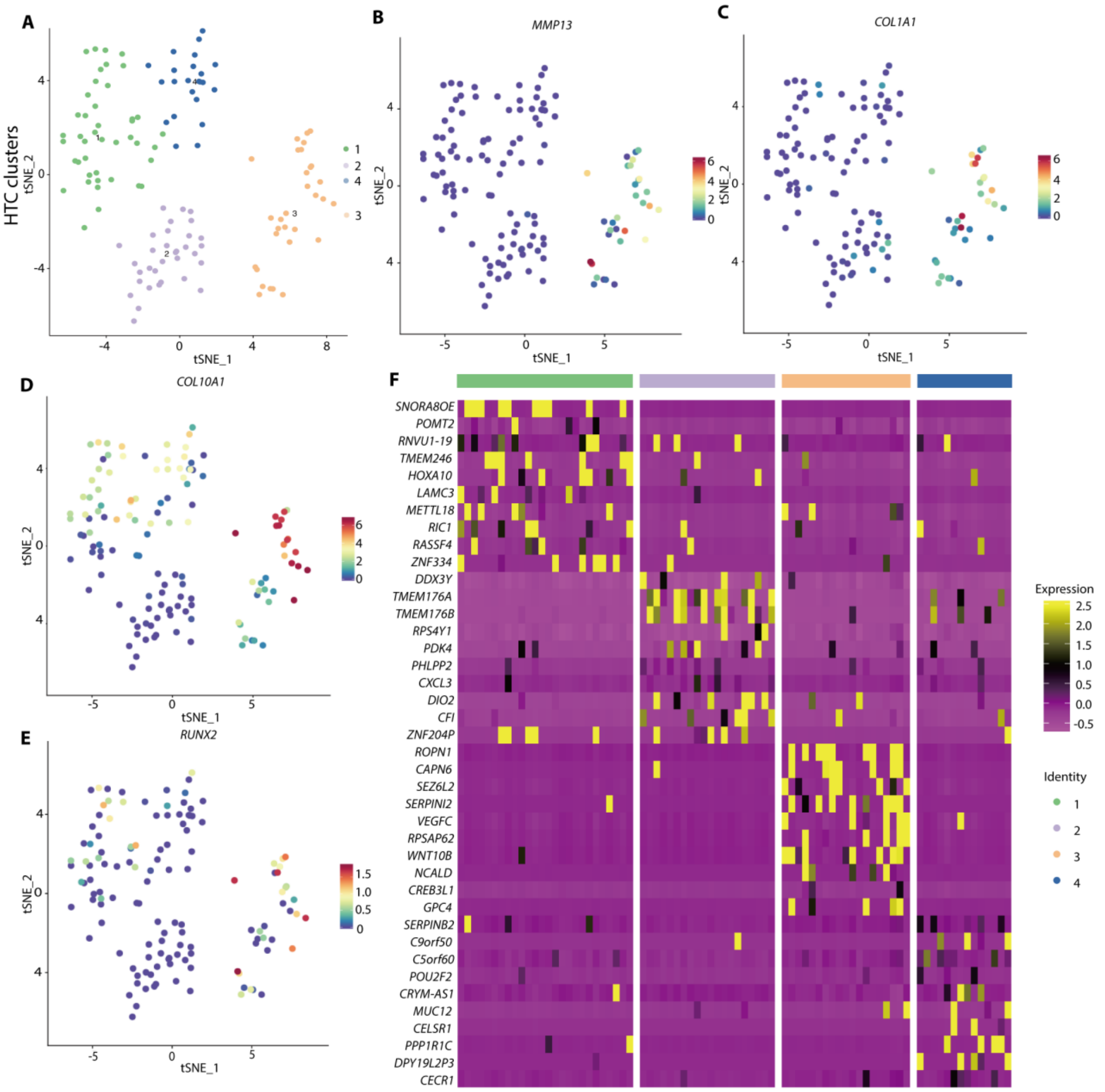


**Fig.S4 HTC subsets definition and characteristic of OA-associated HTC.**

**(A)** Visualization of t-SNE for different subsets of HTCs. **(B-E)** Dot plots showing the expression of *MMP13*, *COL1A1*, *COL10A1* and *RUNX2* of HTC subgroups. **(F)** Heatmap of top 10 different expression genes for each cluster in HTCs.

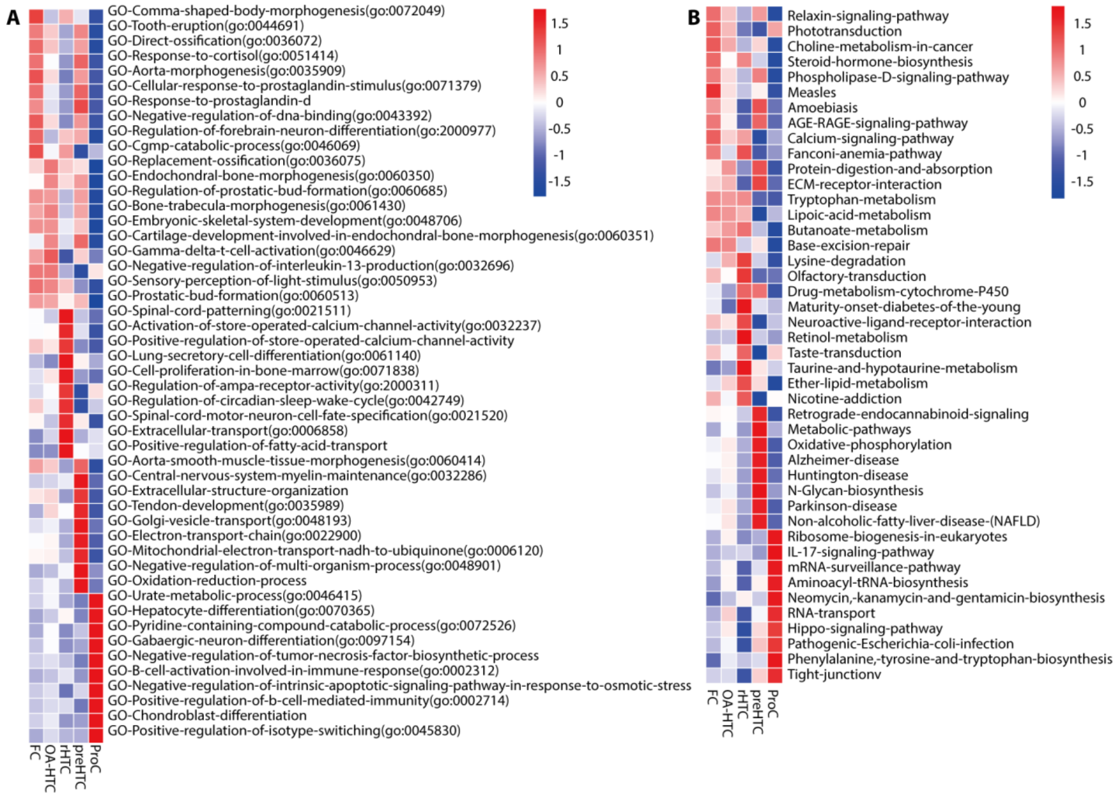


**Fig.S5 KEGG and GO analysis of ProC, PreHTC, rHTC, OA-HTC and FC.**

**(A)** Top 10 GO analysis result of FC, OA-HTC, rHTC, preHTC, and ProC.

**(B)** Top 10 KEGG pathway analysis result of FC, OA-HTC, rHTC, preHTC, and ProC.

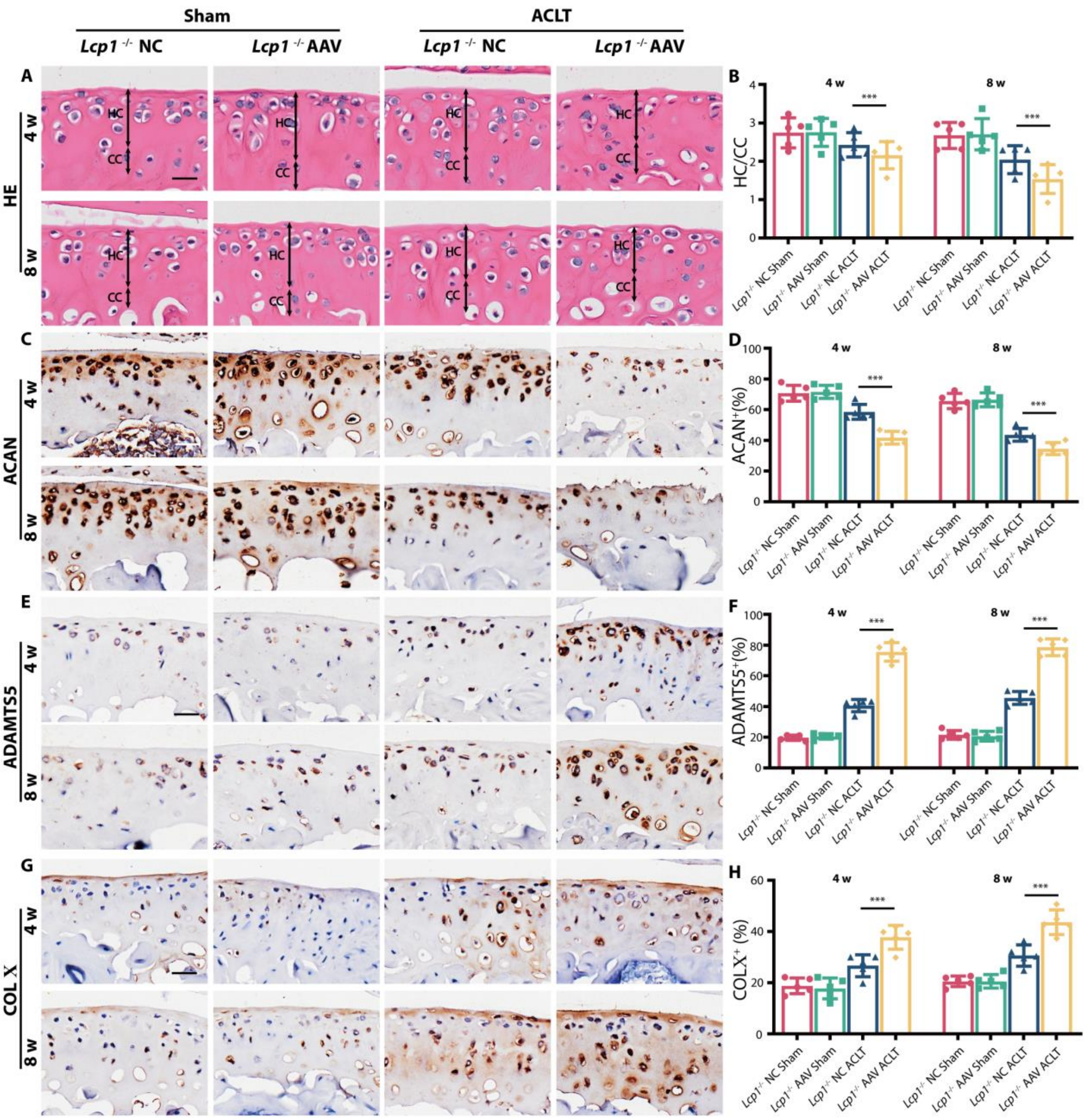


**Fig.S6 Knockdown *Hif-1a* abolishes cartilage protective effects in *Lcp1*$^{-/-}$ mice.**

**(A)** Representative images of Hematoxylin and eosin staining of proximal tibia articular cartilage of *Lcp1*$^{-/-}$ mice with *Hif-1a* AAV or negative control at 4 and 8 weeks after ACLT. Double-headed arrows label range of HC and CC. Scale bar, 50μm. **(B)** Quantitative analysis of HC and CC thickness Ratio. **(C)** Representative images of ACAN protein immunohistochemistry in tibial articular cartilage of *Lcp1*$^{-/-}$ mice with *Hif-1a* AAV or negative control at 4 and 8 weeks after ACLT. Scale bar, 20μm. **(D)** Quantitative analysis of ACAN protein positive area in articular cartilage. **(E)** Representative images of ADAMTS5 protein immunohistochemistry in tibial articular cartilage of

*Lcp1*$^{-/-}$ mice with *Hif-1a* AAV or negative control at 4 and 8 weeks after ACLT. Scale bar, 20μm. **(F)** Quantitative analysis of ADAMTS5 protein positive area in articular cartilage. **(G)** Representative images of COL X protein immunohistochemistry in tibial articular cartilage of *Lcp1*$^{-/-}$ mice with *Hif-1a* AAV or negative control at 4 and 8 weeks after ACLT. Scale bar, 20μm. **(H)** Quantitative analysis of COL X protein positive area in articular cartilage.

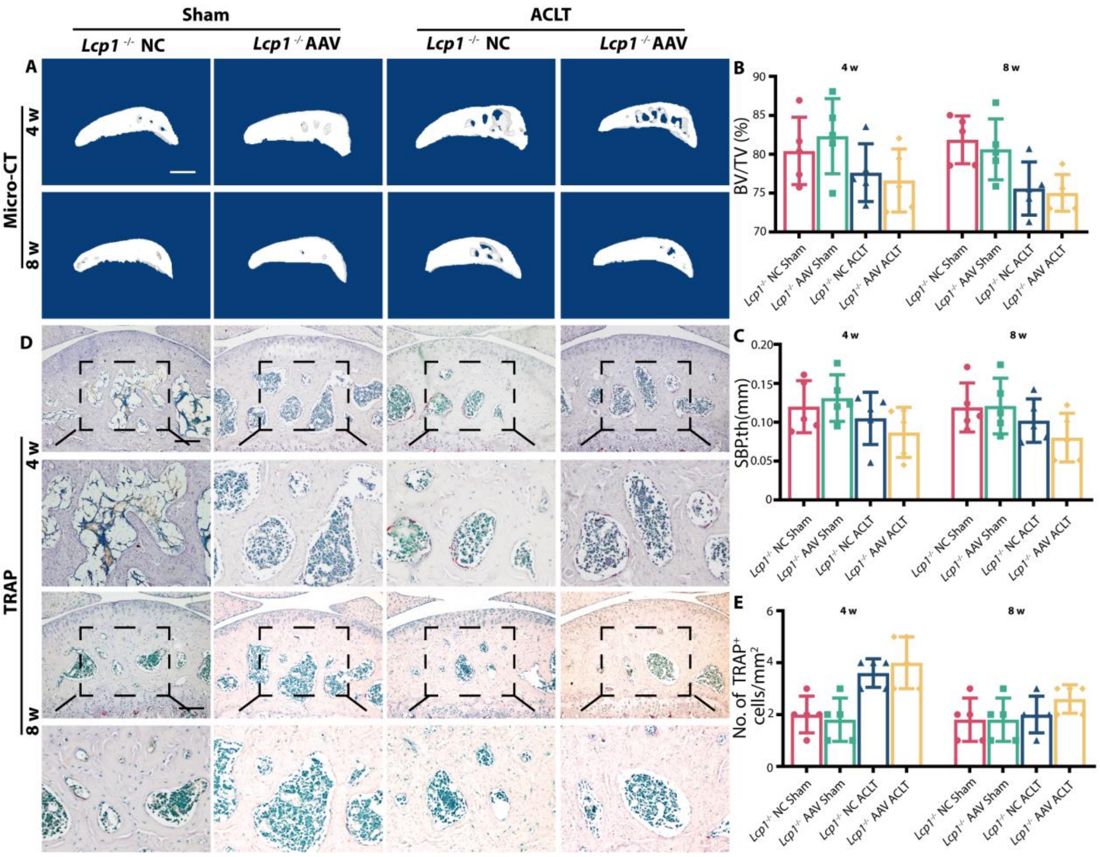


**Fig.S7 Knockdown *Hif-1a* in *Lcp1*$^{-/-}$ mice shows no effects on subchondral bone.**

**(A)** Representative Micro-CT 3D images of tibia subchondral bone of *Lcp1*$^{-/-}$ mice with *Hif-1a* AAV or negative control at 4 and 8 weeks after ACLT. Scale bar, 500μm. **(B-C)** Micro-CT quantitative analysis of tibial subchondral bone, bone volume/tissue volume (BV/TV, %) (B) and subchondral bone plate thickness (SBP. Th, μm) (C). **(D)** TRAP staining image of tibial subchondral bone of *Lcp1*$^{-/-}$ mice with *Hif-1a* AAV or negative control at 4 and 8 weeks after ACLT. Scale bar, 100μm (1,3 raw), 50 μm (2,4 raw). **(E)** Quantitative analysis of TRAP-positive cells in subchondral bone marrow.

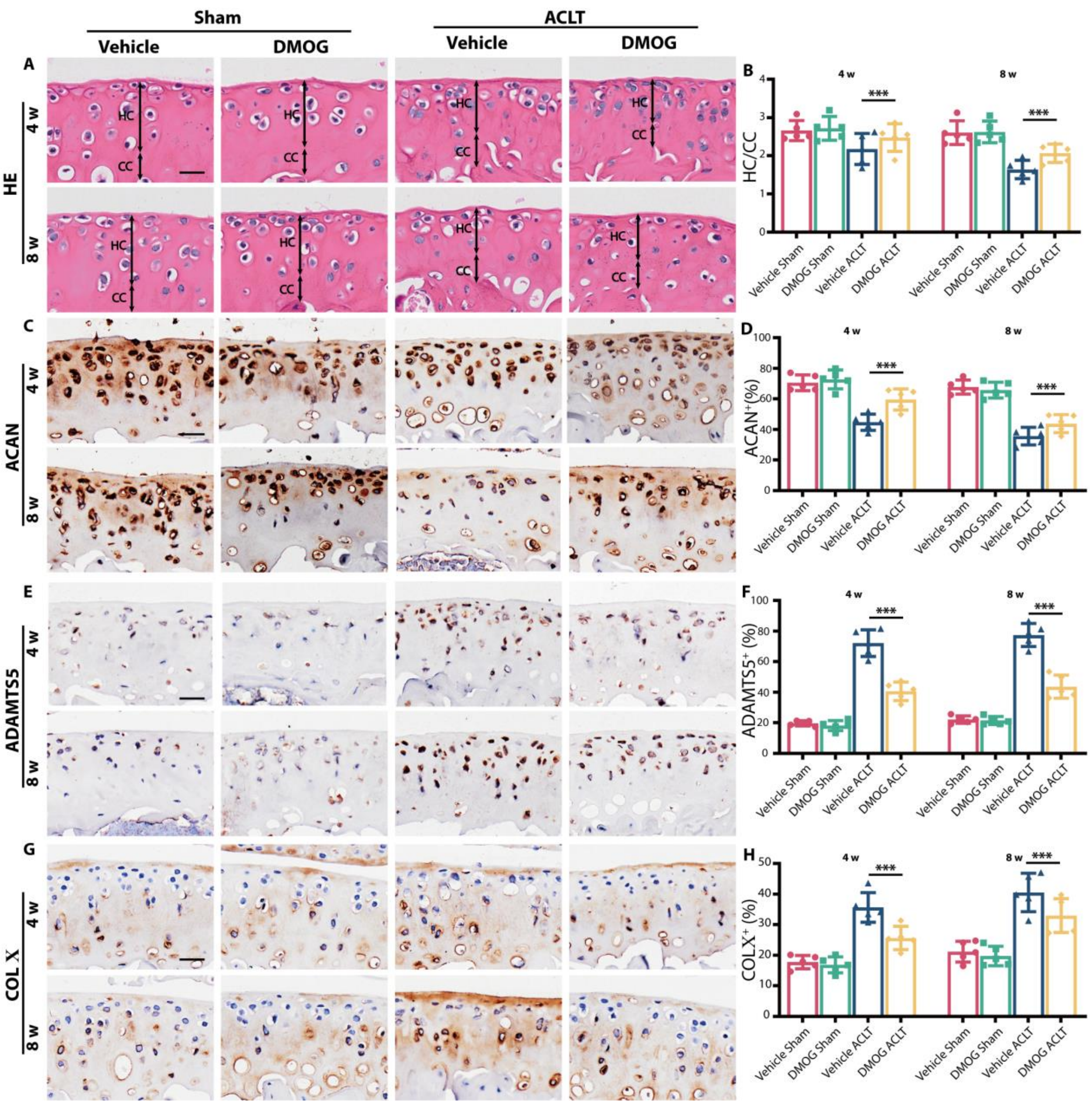


**Fig.S8 DMOG prevents cartilage degeneration.**

**(A)** Representative images of Hematoxylin and eosin staining of proximal tibia articular cartilage of WT mice with DMOG or normal saline at 4 and 8 weeks after ACLT. Double-headed arrows label range of HC and CC. Scale bar, 50μm. **(B)** Quantitative analysis of HC and CC thickness Ratio. **(C)** Representative images of ACAN protein immunohistochemistry in tibial articular cartilage of WT mice with DMOG or normal saline at 4 and 8 weeks after ACLT. Scale bar, 20μm. **(D)** Quantitative analysis of ACAN protein positive area in articular cartilage. **(E)** Representative images of ADAMTS5 protein immunohistochemistry in tibial articular cartilage of WT mice with DMOG or normal saline at 4 and 8 weeks after ACLT. Scale bar, 20μm. **(F)** Quantitative analysis of ADAMTS5 protein positive area in articular cartilage. **(G)** Representative images of COL X protein immunohistochemistry in tibial articular cartilage of WT mice with DMOG or normal saline at 4 and 8 weeks after ACLT. Scale bar, 20μm. **(H)** Quantitative analysis of COL X protein positive area in articular cartilage.

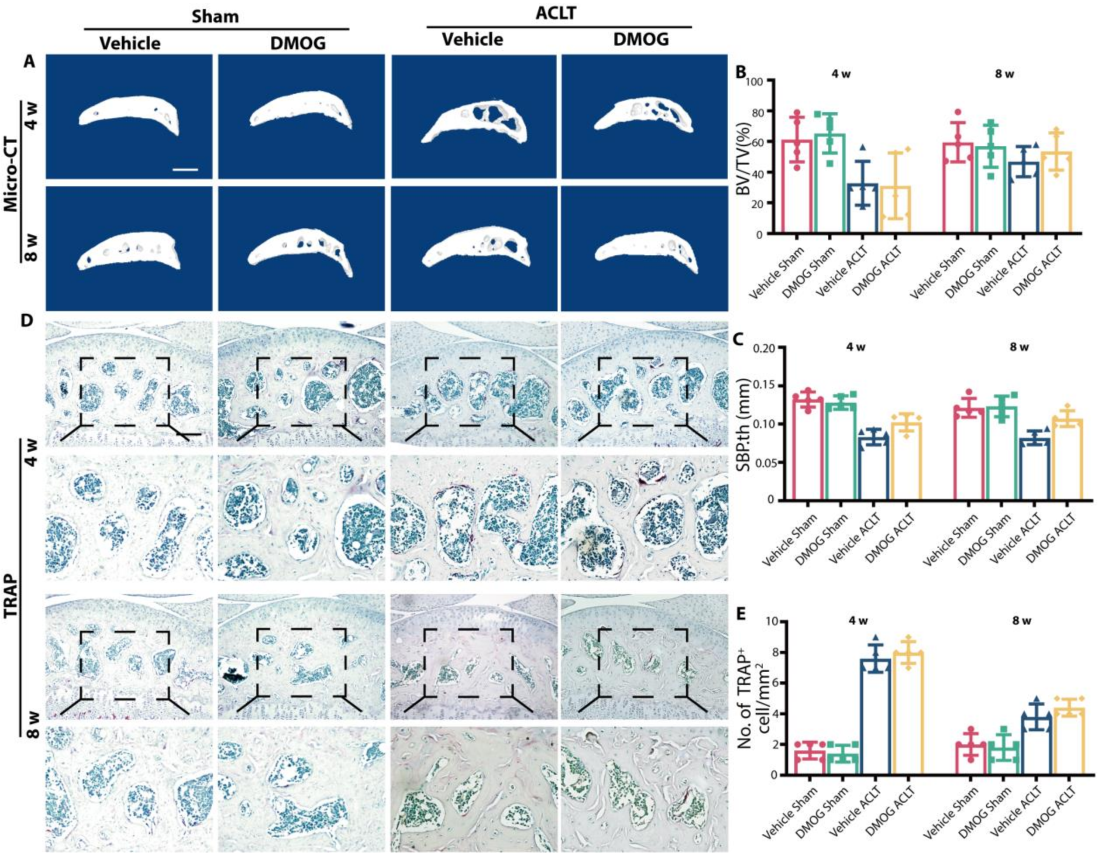


**Fig.S9 DMOG does not affect subchondral bone in OA.**

**(A)** Representative micro-CT 3D images of tibia subchondral bone of WT mice with DMOG or normal saline at 4 and 8 weeks after ACLT. Scale bar, 500μm. **(B-C)** Micro-CT quantitative analysis of tibial subchondral bone, bone volume/tissue volume (BV/TV, %) (B) and subchondral bone plate thickness (SBP. th, μm) (C). **(D)**TRAP staining image of tibial subchondral bone of WT mice with DMOG or normal saline at 4 and 8 weeks after ACLT. Scale bar, 100μm (1,3 raw), 50 μm (2,4 raw). **(E)** Quantitative analysis of TRAP-positive cells in subchondral bone marrow.

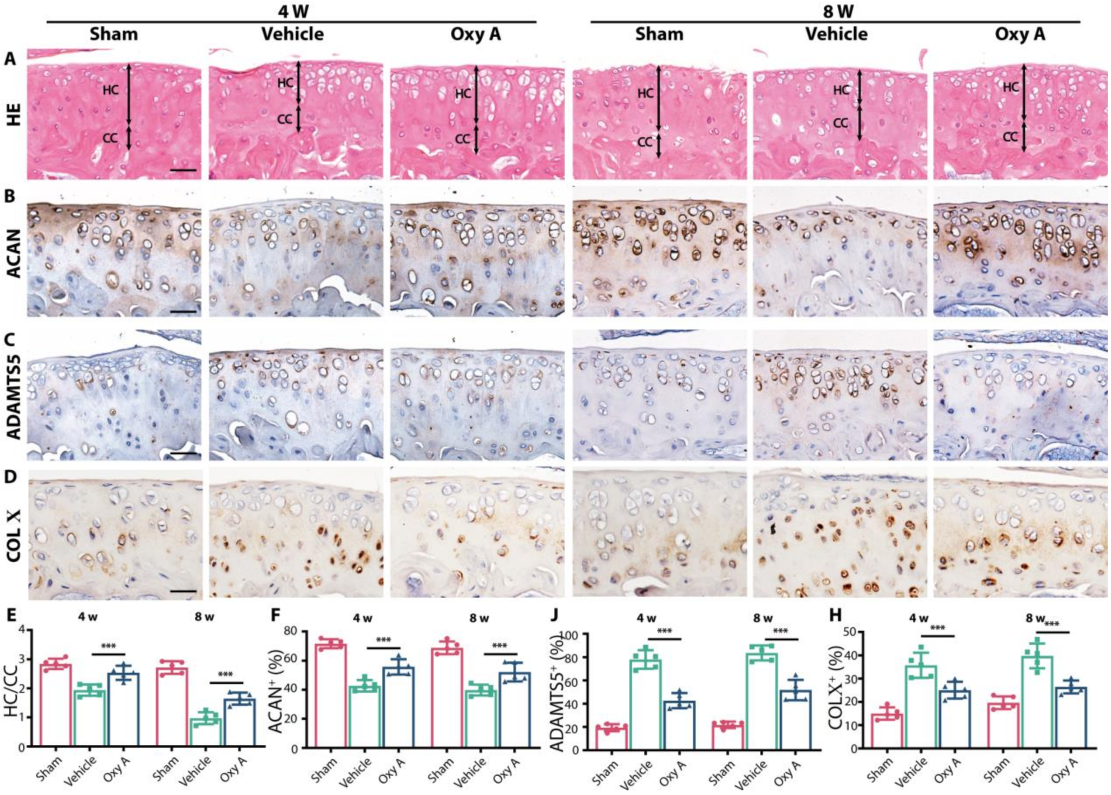


**Fig.S10 Oroxylin A prevents cartilage degeneration.**

**(A)** Representative images of Hematoxylin and eosin staining of proximal tibia articular cartilage of WT mice with Oxy A or normal saline at 4 and 8 weeks after ACLT. Double-headed arrows label range of HC and CC. Scale bar, 50μm. **(B)** Representative images of ACAN protein immunohistochemistry in tibial articular cartilage of WT mice with Oxy A or normal saline at 4 and 8 weeks after ACLT. Scale bar, 20μm. **(C)** Representative images of ADAMTS5 protein immunohistochemistry in tibial articular cartilage of WT mice with Oxy A or normal saline at 4 and 8 weeks after ACLT. Scale bar, 20μm. **(D)** Representative images of COL X protein immunohistochemistry in tibial articular cartilage of WT mice with Oxy A or normal saline at 4 and 8 weeks after ACLT. Scale bar, 20μm. **(E)** Quantitative analysis of HC and CC thickness Ratio. **(F)** Quantitative analysis of ACAN protein positive area in articular cartilage. **(G)** Quantitative analysis of ADAMTS5 protein positive area in articular cartilage. **(H)** Quantitative analysis of COL X protein positive area in articular cartilage. N=5 per group. *P < 0.05, **P < 0.01, and ***P < 0.001.

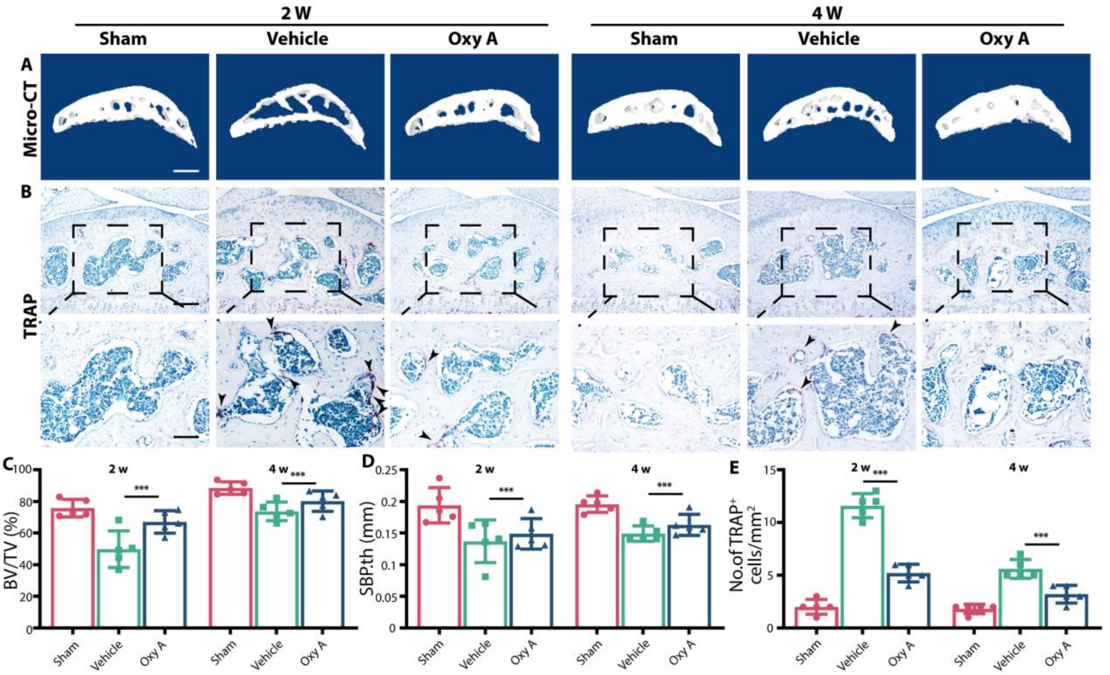


**Fig.S11 Oroxylin A inhibits subchondral bone remodeling via inhibiting osteoclast formation.** **(A)** Representative Micro-CT 3D images of tibia subchondral bone of WT mice with Oxy A or normal saline at 2 and 4 weeks after ACLT. Scale bar, 500μm. **(B)** TRAP staining image of tibial subchondral bone of WT mice with Oxy A or normal saline at 2 and 4 weeks after ACLT. Scale bar,100μm (top). Scale bar, 50μm (bottom). **(C-D)** Micro-CT quantitative analysis of tibial subchondral bone, bone volume/tissue volume (BV/TV, %) (C), and subchondral bone plate thickness (SBP.th, μm) (D). **(E)** Quantitative analysis of TRAP-positive cells in subchondral bone.

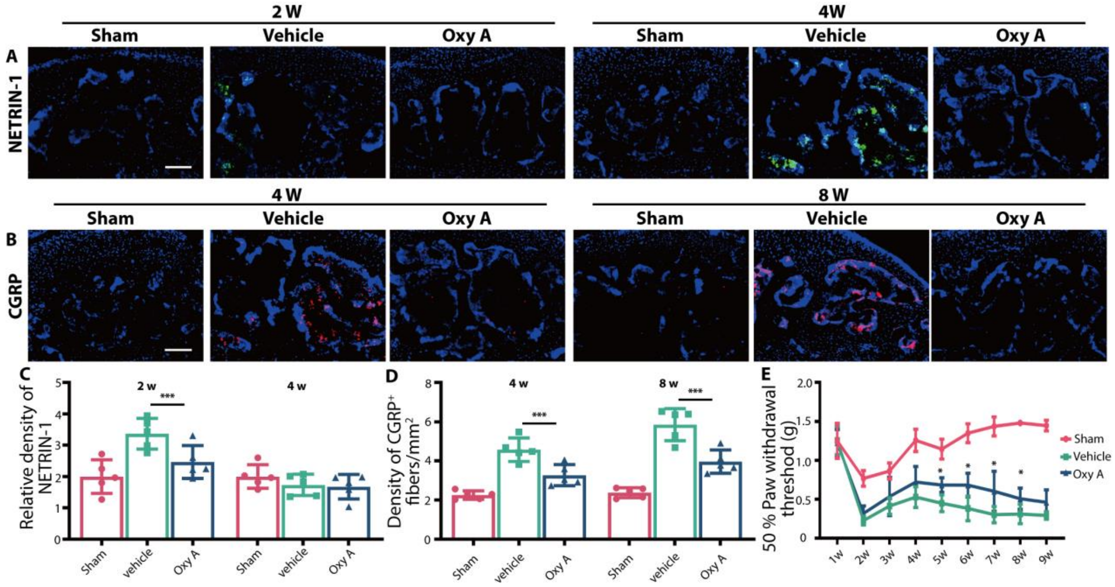


**Fig.S12. Oroxylin A decreases Netrin-1 and CGRP$^{+}$ sensory nerves in the subchondral bone and ameliorates pain in WT mice.**

**(A)** Immunofluorescence staining of NETRIN-1 protein in WT mice with Oxy A or normal saline at 2 and 4 weeks after ACLT. Scale bars, 100 μm. **(B)** Immunofluorescence staining of CGRP$^{+}$ sensory nerve fibers in WT mice with Oxy A or normal saline at 4 and 8 weeks after ACLT surgery. Scale bars, 100 μm. **(C)** Quantitative analysis of density of NERTIN-1 in subchondral bone marrow. **(D)** Quantitative analysis of the density CGRP$^{+}$ nerve fibers in subchondral bone marrow. **(E)** Paw withdrawal threshold was tested at the right hind paw of WT mice with Oxy A or normal saline each week after surgery until 8 weeks. N=5 per group. *$P < 0.05$, **$P < 0.01$, and ***$P < 0.001$.

**Movies S1 Video of $^{18}$F-FMISO PETCT in *Lcp1*$^{-/-}$ mice 4 weeks after ACLT**

**Movies S2 Video of $^{18}$F-FMISO PETCT in WT mice 4 weeks after ACLT**
